# Delayed maturation of hippocampal trisynaptic circuitry underlies the protracted development of episodic-like memory

**DOI:** 10.64898/2026.10.14.759228

**Authors:** Cory McKenzie, Adam I. Ramsaran, Matteo Saderi, Sam Haar, Sheena A. Josselyn, Paul W. Frankland

## Abstract

The ability to form high-resolution episodic memories emerges relatively late in development, but the circuit mechanisms underlying this transition are unknown. CA1 receives entorhinal input through a direct monosynaptic pathway and an indirect trisynaptic pathway via the dentate gyrus and CA3. In mice, CA3→CA1 innervation matured later than MEC→CA1 innervation, with pronounced maturation between P20 and P24. Trisynaptic pathway maturation was necessary for high-resolution memory and was shaped by neuronal activity and early-life experience. Mechanistically, trisynaptic activity promoted perineuronal net assembly around downstream parvalbumin-expressing interneurons, facilitating their maturation and subsequent recruitment during learning to constrain CA1 encoding ensembles. Thus, activity-dependent maturation of trisynaptic circuitry provides a unified circuit and cellular account of a sensitive period for the emergence of episodic-like memory.

## Main Text

The developmental psychologist Jean Piaget proposed that memory comprises two qualitatively distinct systems with different developmental trajectories. He distinguished between memory in the ‘wide sense’, an early-emerging system that supports representations that generalize across experiences, and memory in the ‘strict sense’, which develops later alongside preoperational thought, enabling the conscious recollection of specific past events (*1*). Although Piaget’s terminology predates modern cognitive neuroscience, this distinction closely parallels contemporary accounts separating low-resolution, generalized memory representations—which support statistical learning, semantic knowledge, and category learning—from high-resolution episodic memory, which supports the retrieval of unique experiences.

Developmental studies in humans have largely supported these differential trajectories. Infants exhibit robust statistical learning and category learning from the first months of life (*2–4*), indicating that generalized forms of memory are functional soon after birth. In contrast, behavioral correlates of episodic memory emerge years later (*5–7*). For example, lure discrimination—the ability to distinguish previously encountered items from highly similar lures, a hallmark of high-resolution episodic memory—shows marked development between approximately 4 and 6 years of age and continues to improve into adulthood (*8–10*). Parallel findings have been reported in rodents, where memory expression shifts during the fourth postnatal week from low-resolution, generalized memory to high-resolution, context-specific memory (*11–15*), suggesting that this developmental transition is conserved across species (*6*).

A leading circuit model proposes that these distinct forms of memory arise from parallel pathways within the hippocampus (*16*). The CA1 region receives convergent input from the entorhinal cortex via two anatomically distinct routes: a direct monosynaptic pathway (MSP) from entorhinal cortex to CA1 and an indirect trisynaptic pathway (TSP) through the dentate gyrus (DG) and CA3 before reaching CA1. In the mature hippocampus, the TSP is thought to support high-resolution episodic memory by performing computations such as pattern separation (*17–20*), whereas the MSP preferentially supports generalized, gist-like representations (*16, 21– 23*). Recent studies in adult mice provide causal evidence for this functional dissociation, demonstrating that CA3 inputs to CA1 are required for the formation of high-dimensional memory representations and context-specific memory (*24*, *25*).

If these parallel hippocampal pathways support memories of different resolution, their differential maturation could provide a circuit mechanism for the developmental transition from generalized to high-resolution memory (*10*, *26*). Several lines of evidence suggest that the MSP and TSP indeed follow distinct developmental trajectories. Anatomical studies in non-human primates have reported a prolonged increase in DG and CA3 neuronal populations during postnatal development (*27*), while tracing and electrophysiological studies in perinatal rodents indicate that the MSP begins maturing earlier than the TSP (*28*, *29*). These findings raise the possibility that delayed TSP maturation underlies the late emergence of high-resolution episodic memory, whereas the earlier-maturing MSP is sufficient to support generalized memory.

Here, we addressed this question by combining behavioral analyses, viral circuit tracing, and pathway- and cell-type-specific chemo- and optogenetic manipulations in developing mice. Using contextual fear conditioning, we tracked developmental changes in memory resolution by quantifying the transition from generalized to context-specific memory. We then examined the structural maturation of MEC→CA1 and CA3→CA1 inputs across postnatal development and tested their causal contributions to memory resolution. Our experiments indicate that the TSP anatomically and functionally matures later than the MSP and that this delayed maturation drives the developmental emergence of high-resolution, context-specific memory. Mechanistically, we show that TSP activity promotes perineuronal net (PNN) assembly around downstream PV interneurons, facilitating their maturation and recruitment to support high-resolution, event-based memory. Activity within the developing TSP therefore acts as an instructive signal through which early-life experiences regulate the developmental program for emergence of high resolution, episodic-like memory.

## TSP maturation coincides with the emergence of high-resolution memory

We first defined the developmental trajectory of memory resolution across development. Mice were trained in contextual fear conditioning on postnatal day (P)20, P21, P22, P23, P24, or P60 (i.e., adulthood) and tested 24 hours later in a similar context (B) and 5 hours later in the training context (A) (**Fig. 1A**). Mice trained at P20 or P21 froze equivalently in the training and similar contexts (**Fig. 1B**), indicating generalized memory. In contrast, mice trained at P22 and older selectively froze in the training context (**Fig. 1B**), indicating context-specific memory. Thus, context discrimination increased sharply between P21 and P22 (**Fig. 1C**), marking the emergence of high-resolution memory (*11*, *30*, *31*). The timing of this developmental transition was comparable in males and females **(Fig. S1A-D**) and was not related to differences in learning during training (**Fig. S1E**). Based on these findings, we focused subsequent experiments on P20, P24, and P60 to capture the developmental transition from low-to high-resolution memory. Using these representative ages, we confirmed that the transition in memory resolution was not influenced by weaning (**Fig. S1F-H**) and could not be explained by age-dependent differences in perceptual discrimination (**Fig. S1I-K**).

**Fig. 1.**
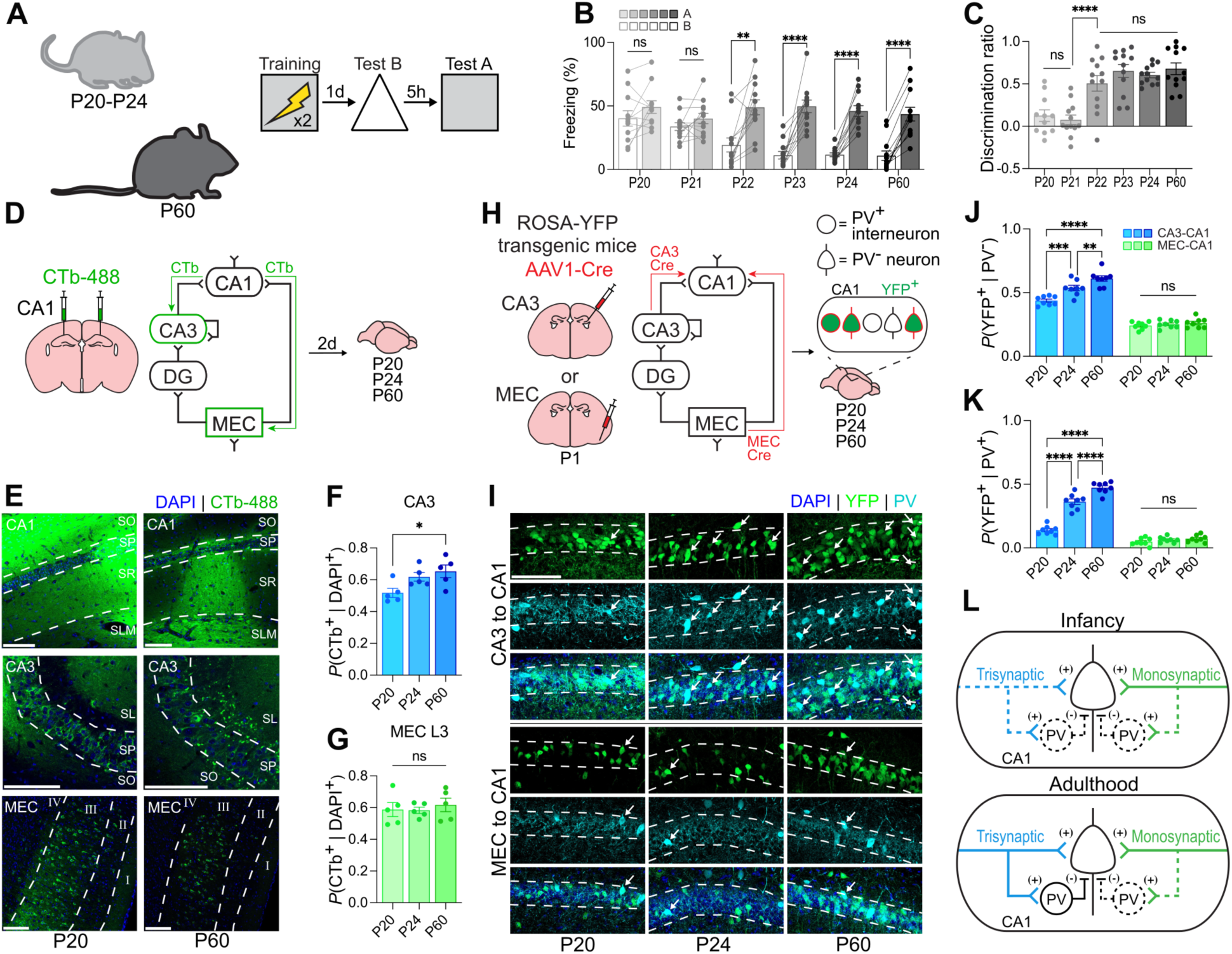
TSP maturation coincides with the emergence of high-resolution memory. **(A)** Contextual fear conditioning across development. Mice were trained at P20–P24 or P60 and tested in a similar context (B) and the training context (A). **(B)** P20 and P21 mice generalized across contexts, whereas mice trained at P22 and older showed context-specific freezing (ANOVA, age × context: *F*_5,65_ = 7.38, *P* < 0.0001). **(C)** Context discrimination increased from P22 (ANOVA, age: *F*_6,68_ = 14.12, *P* < 0.0001). **(D)** Retrograde tracing of CA3→CA1 and MEC→CA1 projections using CTb-488 injected into CA1. **(E)** Representative CTb-488 labeling in CA3 and MEC. **(F and G)** CA3 labeling increased across development (ANOVA, age: *F*_2,12_ = 4.68, *P* = 0.0315), whereas MEC labeling was unchanged (*F*_2,12_ = 0.24, *P* = 0.7872). **(H)** Anterograde transsynaptic tracing of CA3→CA1 and MEC→CA1 projections using AAV1-Cre **in** ROSA-YFP mice. **(I)** Representative transsynaptic YFP labeling in CA1. Arrows indicate co-labeled cells. **(J)** CA3→CA1 labeling increased across development, particularly between P20 and P24, whereas MEC→CA1 labeling was unchanged (ANOVA, age × projection: *F*_2,42_ = 12.15, *P* < 0.0001). **(K)** The developmental increase in CA3→CA1 labeling was greater in PV^+^ interneurons, whereas MEC→CA1 labeling of PV^+^ interneurons remained low (ANOVA, age × projection: *F*_2,42_ = 89.98, *P* < 0.0001). **(L)** Model of earlier maturation of the MSP than the TSP, with delayed development of TSP input to CA1 PV^+^ interneurons. Dashed lines indicate connections which are negligible, while solid lines indicate substantial connections. (+) indicated excitatory synapses while (-) indicated inhibitory synapses. Data are individual mice, mean ± s.e.m. Scale bars: white = 100 µm. \**P* < 0.05; ** *P* < 0.01; \*\*\**P* < 0.001; \*\*\*\**P* < 0.0001.

Because both high- and low-resolution contextual fear memories depend critically on the CA1 region of the hippocampus (*11*, *25*), we next asked whether the developmental emergence of high-resolution memory coincides with maturation of the pathways that converge onto CA1. Given evidence that the TSP matures later than the MSP (*26*, *28*), we used complementary retrograde and anterograde viral tracing strategies to examine the development of these inputs. We first injected the retrograde tracer cholera toxin subunit B (CTb-488) into CA1 and brains were analyzed on P20, P24, or P60. Labeled neurons in CA3 and medial entorhinal cortex (MEC) layer 3 were quantified (**Fig. 1D**). Robust retrograde labeling was observed in both regions at all ages, confirming successful labeling of CA3→CA1 and MEC→CA1 projections (**Fig. 1E**). The proportion of labeled DAPI-normalized CTb-labeled neurons in CA3 increased progressively from P20 to adulthood (**Fig. 1F**), consistent with prolonged structural maturation of the TSP. In contrast, the proportion of labeled neurons in MEC was similar across ages, suggesting that MEC→CA1 inputs undergo little further structural change after P20 (**Fig. 1G**). As expected, retrograde tracing from DG and CA1 revealed distinct layer-specific labeling patterns in MEC, consistent with the known organization of TSP-versus MSP-projecting neurons in entorhinal cortex (**Fig. S2A-E**).

To identify the postsynaptic targets of these projections, we next used a complementary anterograde transsynaptic tracing approach. AAV1 expressing Cre recombinase (AAV1-Cre) was injected into either CA3 or MEC of Cre-dependent ROSA-YFP reporter mice and brains were analyzed on P20, P24, or P60. YFP expression at the injection site was comparable across ages, and control experiments confirmed that transsynaptic labeling was restricted to monosynaptically connected neurons (**Fig. S2F-M**). Transsynaptic transfer of Cre from transfected presynaptic neurons resulted in YFP labeling of CA1 neurons (**Fig. 1H**). Robust transsynaptic labeling in CA1 was observed following injections into both CA3 and MEC at all ages (**Fig. 1I**). To determine whether these pathways differentially target inhibitory microcircuits, tissue was counterstained with antibodies for parvalbumin (PV) or somatostatin (SST), and the proportions of transsynaptically labeled interneurons were quantified.

Consistent with the retrograde tracing experiments, transsynaptic labeling following MEC injections was comparable across ages, suggesting little further structural maturation of MEC→CA1 inputs after P20. In contrast, transsynaptic labeling of CA1 neurons following CA3 injections increased markedly between P20 and adulthood, with the largest increase occurring between P20 and P24 (**Fig. 1J**). This developmental increase was evident in both PV^-^ (i.e., putative pyramidal neurons) and PV^+^ neurons but was substantially greater in PV^+^ neurons (**Fig. 1K**). By contrast, we observed no developmental increase in transsynaptic labeling of SST^+^ neurons (**Fig. S3A-C**). Thus, the developmental increase in CA3→CA1 innervation of CA1 PV interneurons was substantially greater than that observed for MEC→CA1 inputs and was not a general feature of inhibitory interneurons. Because CA1 PV interneurons are critical for generating sparse, high-resolution hippocampal representations through feedforward inhibition (*24*, *32–35*), these findings suggest that the increased recruitment of CA1 PV interneurons during the fourth postnatal week (*11*) may depend on maturation of the TSP (**Fig. 1L**).

## TSP maturation enables recruitment of CA1 PV interneurons

To directly address this possibility, we expressed a blue-light-sensitive excitatory opsin in either CA3 or MEC, photostimulated the transfected neurons *in vivo*, and quantified the resulting c-Fos induction in PV^+^ and PV^-^ CA1 neurons of P20, P24, and P60 mice (**Fig. 2A**). Photostimulation of CA3 increased c-Fos expression in PV^-^ CA1 neurons at all ages examined, whereas photostimulation of MEC had little effect (**Fig. 2C**). Baseline c-Fos expression in PV^-^ neurons was elevated at P20 (**Fig. 2C**), consistent with previous observations of heightened network activity during early postnatal development (*11*, *36*). A different pattern was observed in PV^+^ interneurons. CA3 stimulation failed to recruit PV^+^ interneurons at P20 but produced robust c-Fos expression at P24 and P60, whereas MEC stimulation had no significant effect at any age (**Fig. 2D**). This age-dependent increase in interneuron recruitment was specific to PV interneurons, as neither CA3 nor MEC stimulation significantly recruited SST interneurons at any age (**Fig. S3D-F**).

**Fig. 2.**
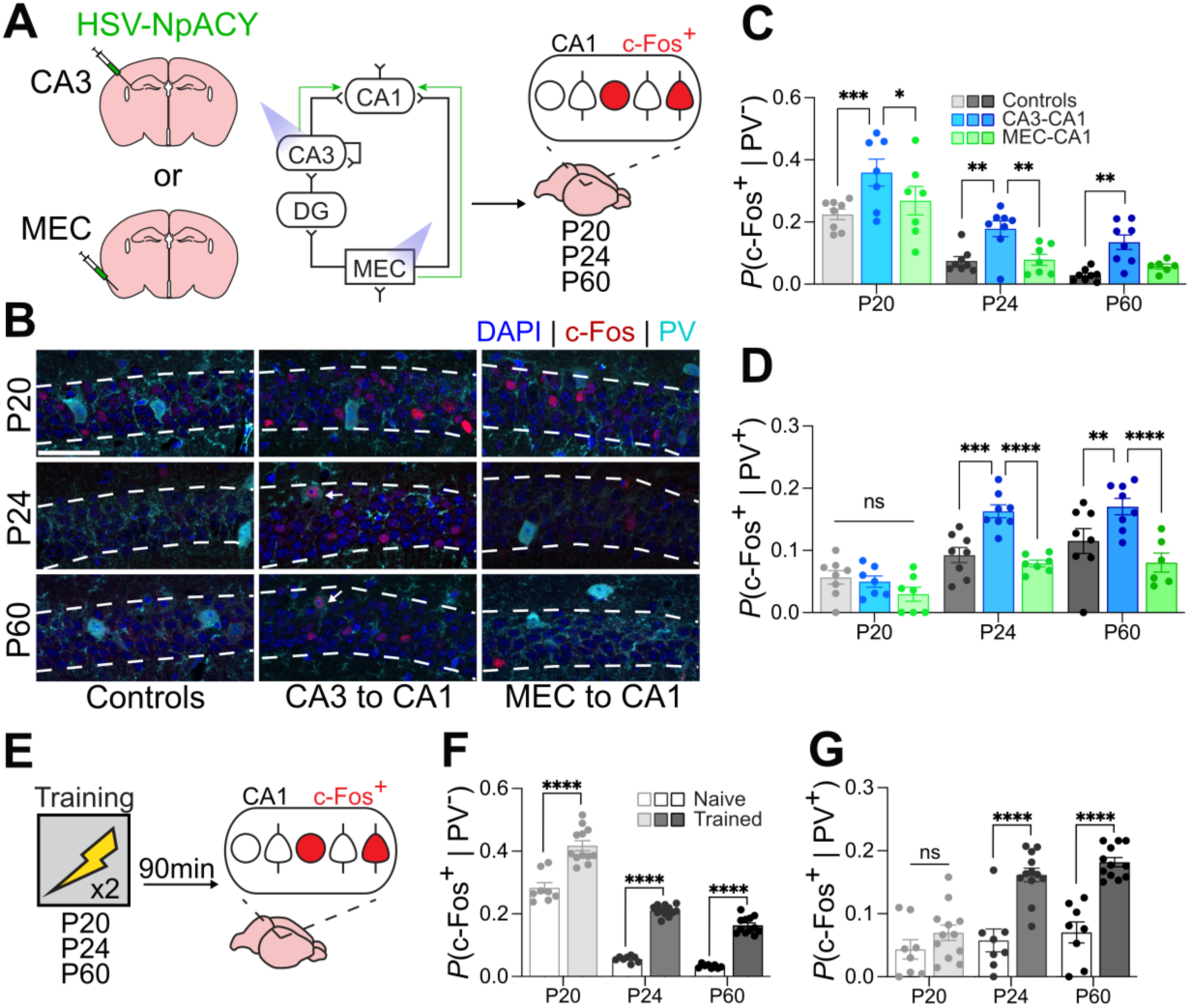
TSP maturation enables recruitment of CA1 PV interneurons. **(A)** Channelrhodopsin-2 was expressed in CA3 or MEC using HSV-NpACY, followed by photostimulation and quantification of c-Fos expression in PV-positive and PV-negative CA1 neurons at P20, P24, and P60. **(B)** Representative c-Fos and PV labeling in CA1 following CA3 or MEC stimulation**. A**rrows indicate co-labeled cells. **(C)** CA3 stimulation increased c-Fos expression in PV**^-^** CA1 neurons at all ages, whereas MEC stimulation had no effect (ANOVA, age × projection: *F*_4,58_ = 0.20, *P* = 0.9385). **(D)** CA3 stimulation recruited PV^+^ interneurons at P24 and P60, but not P20, whereas MEC stimulation did not recruit PV^+^ interneurons at any age (ANOVA, age × projection: *F*_4,58_ = 3.27, *P* = 0.0173). **(E)** c-Fos expression was quantified following contextual fear conditioning or in naïve controls. **(F)** Learning increased c-Fos expression in PV**^-^** CA1 neurons at all ages (ANOVA, age × training: *F*_2,54_ = 0.71, *P* = 0.4956). **(G)** Learning-induced recruitment of PV^+^ interneurons emerged between P20 and P24 (ANOVA, age × training: *F*_2,54_ = 6.43, *P* = 0.0031). Data are individual mice, mean ± s.e.m, Scale bars: white = 100 µm. \**P* < 0.05; ** *P* < 0.01; \*\*\**P* < 0.001; \*\*\*\**P* < 0.0001.

Having established that CA3→CA1 inputs acquire the capacity to recruit CA1 PV interneurons during development, we examined whether a corresponding transition occurs following learning. Mice underwent contextual fear conditioning or remained in their home cage before c-Fos expression was quantified in CA1 (**Fig. 2E**). In PV^-^ neurons, learning increased c-Fos expression at all ages, although baseline activity was again elevated at P20 and induction was more pronounced at P24 and P60 (**Fig. 2F**). In contrast, learning failed to recruit PV^+^ neurons at P20 but robustly increased c-Fos expression in PV^+^ neurons at P24 and P60 (**Fig. 2G**). Thus, the developmental increase in CA3→CA1 innervation of PV interneurons is accompanied by the emergence of PV interneuron recruitment following both artificial, selective pathway activation and memory formation. These findings identify the developmental recruitment of CA1 PV interneurons by TSP as a candidate circuit mechanism for the emergence of high-resolution memory.

## TSP-dependent PV recruitment determines memory resolution

We next asked whether the TSP is required to recruit CA1 PV interneurons during learning. We expressed a red-light-sensitive inhibitory opsin in either CA3 or MEC and implanted optic fibers above CA1 to inhibit CA3→CA1 or MEC→CA1 terminals during contextual fear conditioning (**Fig. 3A**). We then quantified learning-induced c-Fos expression in PV^+^ and PV^-^ CA1 neurons (**Fig. 3B**). At P20, learning induced robust c-Fos expression in PV^-^ neurons, which was moderately reduced by inhibition of CA3→CA1, but not MEC→CA1, projections (**Fig. 3C**). Learning failed to recruit PV^+^ neurons at this age regardless of pathway inhibition (**Fig. 3D**). In contrast, at P24, inhibition of CA3→CA1, but not MEC→CA1, projections increased c-Fos expression in PV^-^ neurons while markedly reducing recruitment of PV^+^ interneurons (**Fig. 3E-F**). Thus, maturation of the TSP enables recruitment of CA1 PV interneurons during learning, which constrains the size of the CA1 encoding ensemble.

**Fig. 3.**
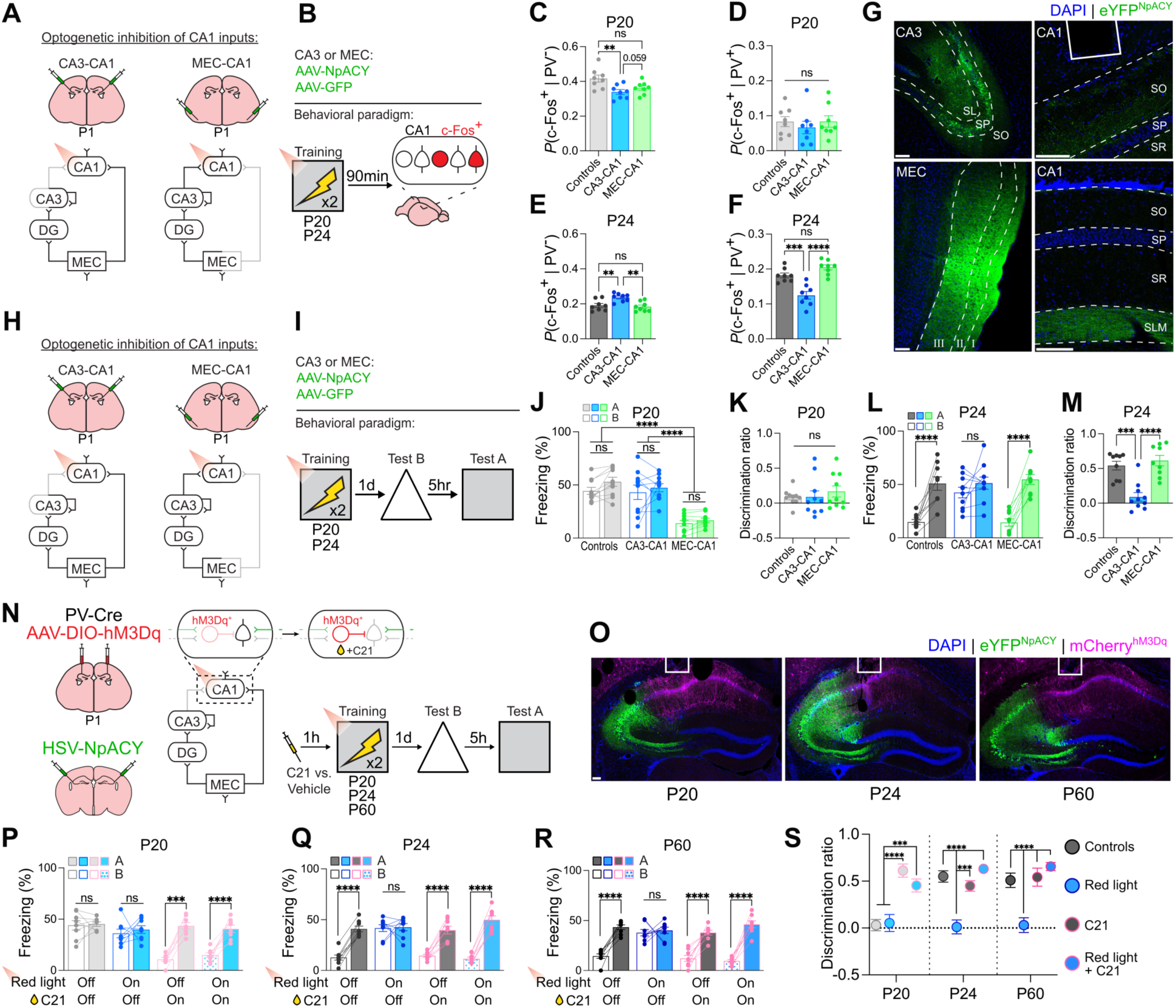
TSP-dependent PV interneuron recruitment determines memory resolution. (A and. **B)** CA3→CA1 or MEC→CA1 terminals were inhibited during contextual fear conditioning before quantification of c-Fos expression in CA1. **(C and D)** At P20, CA3→CA1 inhibition reduced c-Fos expression in PV**^-^** neurons (ANOVA, projection: *F*_2,21_ = 6.03, *P* = 0.0085), whereas PV^+^ interneurons were not recruited under any condition (ANOVA, projection: *F*_2,21_ = 0.36, *P* = 0.3645). **(E and F)** At P24, CA3→CA1, but not MEC→CA1, inhibition increased c-Fos expression in PV**^-^** neurons (ANOVA, projection: *F*_2,21_ = 8.61, *P* = 0.0019) and reduced PV^+^ interneuron recruitment (ANOVA, projection: *F*_2,21_ = 23.73, *P* < 0.0001). **(G)** Representative AAV-NpACY expression in CA3 and MEC and corresponding terminals in CA1. **(H and I)** CA3→CA1 or MEC→CA1 terminals were inhibited during training before memory testing. **(J and K)** At P20, CA3→CA1 inhibition did not alter subsequent freezing, whereas MEC→CA1 inhibition reduced freezing (ANOVA, projection: *F*_2,27_ = 28.40, *P* < 0.0001). Discrimination was comparable across groups (ANOVA, projection: *F*_2,27_ = 0.43, *P* = 0.6533). **(L and M)** At P24, CA3→CA1, but not MEC→CA1, inhibition caused mice to generalize across contexts (ANOVA, projection × context: *F*_2,25_ = 8.10, *P* = 0.0019), reducing context discrimination (ANOVA, projection: *F*_2,25_ = 18.70, *P* < 0.0001). **(N)** Combined CA3→CA1 inhibition and chemogenetic activation of CA1 PV^+^ interneurons during contextual fear conditioning. **(O)** Representative HSV-NpACY expression in CA3 and AAV-DIO-hM3Dq expression in CA1 PV^+^ interneurons. **(P)** At P20, PV^+^ interneuron activation induced context-specific freezing despite CA3→CA1 inhibition (ANOVA, injection × context: *F*_1,28_ = 27.08, *P* < 0.0001; photostimulation × injection × context: *F*_1,28_ = 0.83, *P* = 0.3693). **(Q and R)** At P24 and P60, CA3→CA1 inhibition caused mice to generalize across contexts, whereas concurrent PV^+^ interneuron activation restored context-specific freezing (P24: ANOVA, photostimulation × injection × context: *F*_1,28_ = 25.22, *P* < 0.0001; P60: *F*_1,28_ = 17.75, *P* = 0.0002). **(S)** PV^+^ interneuron activation restored context discrimination following CA3→CA1 inhibition at all ages (ANOVA, age × injection × photostimulation: *F*_2,84_ = 12.18, *P* < 0.0001). Data are individual mice, mean ± s.e.m. Scale bars: white = 100 µm. \**P* < 0.05; ** *P* < 0.01; \*\*\**P* < 0.001; \*\*\*\**P* < 0.0001.

To determine the behavioral consequences of these pathway-specific recruitment patterns we inhibited either CA3→CA1 or MEC→CA1 terminals during contextual fear conditioning and assessed memory 24 hours later (**Fig. 3H-I**). Because CA3→CA1 projections are still immature at P20, we predicted that contextual fear learning at this age would depend primarily on the MSP. Consistent with previous findings, control mice trained at P20 exhibited generalized freezing across contexts, reflecting low-resolution memory (**Fig. 3J-K**). Silencing CA3→CA1 terminals had no effect on subsequent freezing (**Fig. 3J**), suggesting that the immature TSP makes little contribution to learning at this age. In contrast, silencing MEC→CA1 terminals markedly impaired contextual fear acquisition, reducing subsequent freezing (**Fig. 3J**). Together, these findings indicate that contextual fear learning at P20 depends predominantly on the MSP.

We performed the same manipulations in P24 mice. Because CA3→CA1 projections undergo substantial maturation between P20 and P24, we predicted that silencing this pathway during learning would preserve acquisition but reduce memory resolution, since memory could be alternately supported MEC→CA1 projections. Consistent with this prediction, inhibition of CA3→CA1 terminals did not impair contextual fear acquisition, but mice exhibited robust and equivalent freezing in contexts A and B, indicating generalized memory and reduced memory resolution (**Fig. 3L-M**). In contrast, inhibition of MEC→CA1 terminals had no effect on either learning or memory resolution (**Fig. 3L-M**). Thus, by P24, the TSP is required not for memory formation itself, but does determine the resolution of the resulting memory.

These findings suggested that TSP-dependent recruitment of PV interneurons might be the mechanism through which the pathway promotes memory resolution. If so, directly increasing PV interneuron activity should overcome the effects of an immature or inhibited TSP. To test this, we expressed the excitatory DREADD hM3Dq selectively in CA1 PV^+^ neurons in PV-Cre mice and combined chemogenetic PV^+^ neuronal activation with optogenetic inhibition of CA3→CA1 terminals during contextual fear conditioning (**Fig. 3N**). As expected, P20 mice generalized across contexts, and this phenotype was unaffected by CA3→CA1 inhibition (**Fig. 3P**). In contrast, chemogenetic activation of CA1 PV^+^ interneurons caused mice to selectively freeze in the training context, indicating the precocious emergence of high-resolution memory (**Fig. 3P, S**). We next tested whether PV^+^ neuronal activation could similarly restore context-specific memory following TSP inhibition at P24 and P60. At both ages, inhibition of CA3→CA1 terminals caused mice to generalize across contexts (**Fig. 3Q-R**). Concurrent activation of CA1 PV^+^ neurons restored context-specific freezing despite continued inhibition of CA3→CA1 terminals (**Fig. 3Q-S**). Together, these results support a model in which maturation of the TSP enables CA3→CA1 inputs to recruit CA1 PV interneurons during learning, thereby constraining CA1 encoding ensembles and promoting the formation of high-resolution contextual memories.

## Early-life experience shifts TSP maturation and memory resolution

Because early-life adversity and environmental enrichment have previously been shown to delay or accelerate, respectively, the emergence of high-resolution memory, we hypothesized that these effects are accompanied by corresponding changes in TSP maturation. To test this, we bidirectionally manipulated early-life experience using either a limited bedding and nesting model of early-life adversity (ELA) or an enriched housing model of early-life enrichment (ELE) (*31*, *37*, *38*). Consistent with our previous studies, ELA reduced developmental weight gain, whereas ELE did not; neither manipulation altered the timing of eye-opening (*31*) (**Fig. S4A-B**). Using the same transsynaptic viral tracing approach described above, we injected AAV1-Cre into CA3 at P1 and quantified transsynaptic YFP labeling in CA1 at P20 and P24 (**Fig. 4A**). Early-life experience bidirectionally altered maturation of CA3→CA1 inputs. Relative to controls, ELE increased transsynaptic labeling of both PV^-^ and PV^+^ CA1 neurons at P20, with a particularly pronounced increase in labeled PV^+^ neurons (**Fig. 4B-C**). Conversely, ELA reduced transsynaptic labeling of both populations at P24 (**Fig. 4D-E**). Thus, enrichment and adversity were associated with accelerated and delayed maturation, respectively, of CA3→CA1 inputs.

**Fig. 4.**
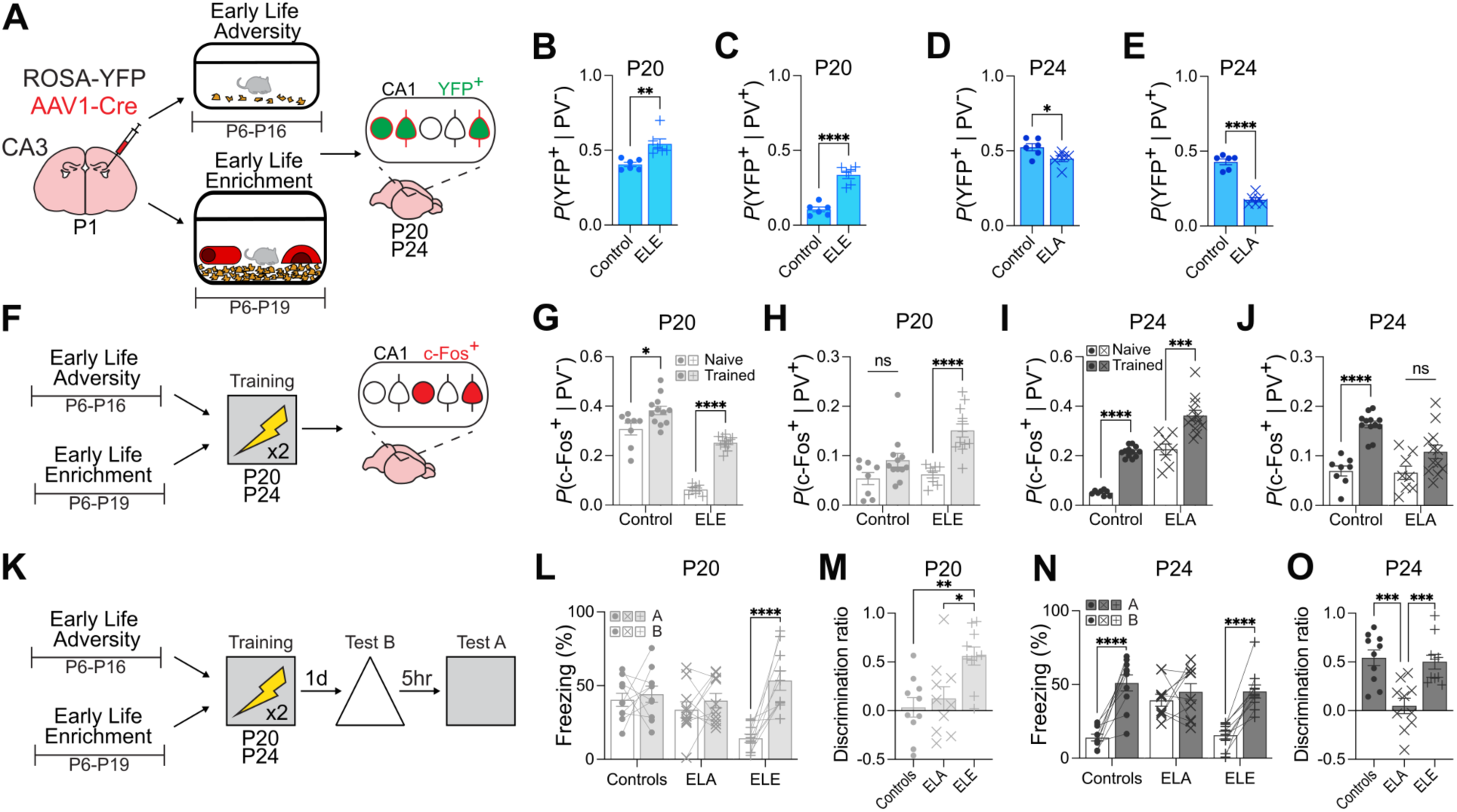
Early-life experiences shift trajectory for TSP maturation and memory resolution. **(A)** Early-life adversity (ELA) and early-life enrichment (ELE) interventions were combined with AAV1-Cre transsynaptic tracing from CA3 to assess CA3→CA1 maturation at P20 and P24. **(B and C)** At P20, ELE increased CA3→CA1 labeling of PV^-^ neurons (unpaired *t*-test, *t*_10_ = 3.79, *P* = 0.0035) and PV^+^ interneurons (unpaired *t*-test, *t*_10_ = 7.76, *P* < 0.0001). **(D and E)** At P24, ELA reduced CA3→CA1 labeling of PV^-^ neurons (unpaired *t*-test, *t*_10_ = 2.43, *P* = 0.0353) and PV^+^ interneurons (unpaired *t*-test, *t*_10_ = 10.45, *P* < 0.0001). **(F)** Learning-induced c-Fos expression was assessed following contextual fear conditioning in P20 ELE and P24 ELA mice. **(G and H)** At P20, ELE decreased c-Fos expression in PV^-^ neurons (ANOVA, experience: *F*_1,36_ = 149.60, *P* < 0.0001) and increased PV^+^ interneuron recruitment (ANOVA, *F*_1,36_ = 6.76, *P* = 0.0134). **(I and J)** At P24, ELA increased c-Fos expression in PV_-_ neurons (ANOVA, *F*_1,36_ = 98.70, *P* < 0.0001) and reduced PV^+^ interneuron recruitment (ANOVA, *F*_1,36_ = 6.28, *P* = 0.0169). **(K)** Memory was assessed following ELA or ELE. **(L and M)** At P20, ELE mice selectively froze in the training context (ANOVA, experience × context: *F*_2,27_ = 6.31, *P* = 0.0056), resulting in increased context discrimination (ANOVA, experience: *F*_2,27_ = 7.14, *P* = 0.0032). **(N and O)** At P24, ELA mice generalized across contexts (ANOVA, experience × context: *F*_2,27_ = 7.63, *P* = 0.0024), resulting in reduced context discrimination (ANOVA, experience: *F*_2,27_ = 12.20, *P* = 0.0002). Data are individual mice, mean ± s.e.m. \**P* < 0.05; ** *P* < 0.01; \*\*\**P* < 0.001; \*\*\*\**P* < 0.0001.

These experience-dependent changes in TSP maturation were accompanied by corresponding changes in CA1 recruitment during learning. We quantified learning-induced c-Fos expression following contextual fear conditioning (**Fig. 4F**). P20 mice raised under ELE exhibited a more mature pattern of CA1 recruitment, with increased recruitment of PV^+^ neurons and reduced activation of PV^-^ neurons (**Fig. 4G-H**). Conversely, P24 mice raised under ELA retained an immature pattern of CA1 recruitment, with reduced PV^+^ neuron recruitment and increased activation of PV^-^ neurons (**Fig. 4I-J**). Together, these findings show that early-life experience bidirectionally shifts TSP maturation, and these shifts parallel changes in CA1 inhibitory recruitment, encoding ensemble size, and the developmental emergence of high-resolution memory.

The anatomical and functional changes induced by early-life experience were also associated with corresponding shifts in memory resolution (**Fig. 4K**). Consistent with previous work (*31*), P20 mice raised under ELE selectively froze selectively in the training context, indicating precocious context-specific memory (**Fig. 4L-M**). In contrast, P24 mice exposed to ELA froze equivalently across contexts, indicating persistent memory generalization (**Fig. 4N-O**). Thus, ELE and ELA bidirectionally shifted the developmental trajectory of memory resolution, and these effects were comparable in males and females (**Fig. S4C-K**). Perineuronal nets (PNNs) are specialized extracellular matrix structures that condense around PV interneurons during development. We previously showed that their formation in CA1 between P20 and P24 is required for PV interneuron-mediated constraint of CA1 encoding ensembles and the emergence of high-resolution memory (*11*). We therefore asked whether early-life experience also alters PNN assembly. ELE increased PNNs at P20, whereas ELA reduced PNNs at P24, with similar effects in trained and naïve mice (**Fig. S5A-C**) Thus, experience-dependent shifts in memory resolution closely paralleled changes in maturation of the TSP and PNN accumulation around CA1 PV interneurons.

## TSP activity instructs PNN maturation during a sensitive period

Maturation of the hippocampus, including PV interneurons, requires neuronal activity that originates in the MEC and propagates through the extended hippocampal circuitry during development (*39*), yet the contribution of TSP versus MSP in this process is unclear. Similar to other systems (*40*), we reasoned that propagating activity along the TSP during the third postnatal week may regulate the timing of its maturation. To test this, we expressed the excitatory opsin CheRiff in CA3 and bidirectionally manipulated activity of CA3→CA1 terminals during development using either high-frequency (50 Hz) or low-frequency (1 Hz) photostimulation (**Fig. 5A**). We first combined these manipulations with AAV1-Cre transsynaptic tracing to examine structural maturation of the pathway. High-frequency stimulation of CA3→CA1 increased transsynaptic labeling of both PV^-^ and PV^+^ CA1 neurons at P20 (**Fig. 5B-C**), whereas low-frequency stimulation reduced transsynaptic labeling at P24 (**Fig. 5D-E**). Thus, activity within the developing TSP bidirectionally regulates its structural maturation.

**Fig. 5.**
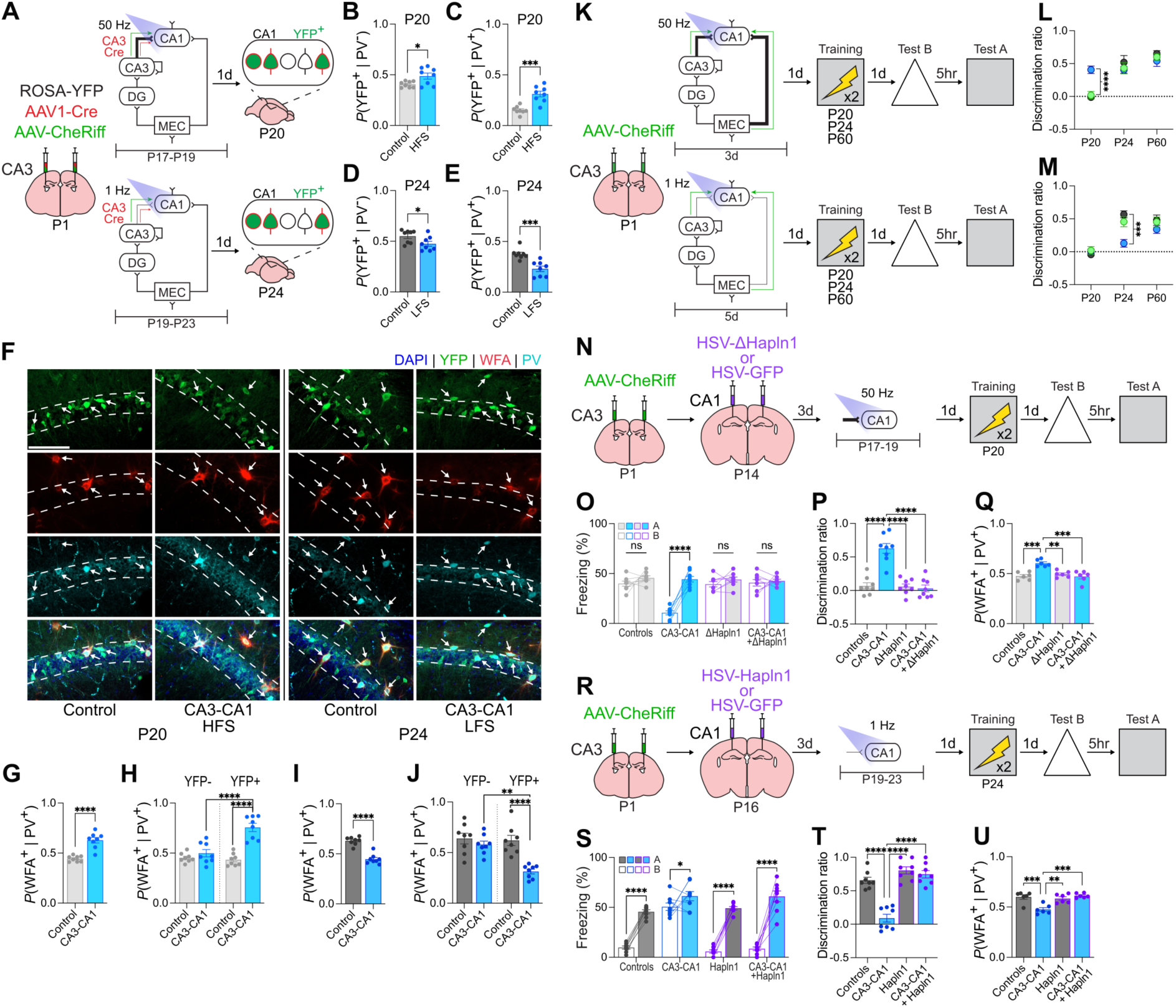
TSP activity instructs PNN maturation during a sensitive period. **(A)** TSP activity was increased or reduced during development using high- or low-frequency CA3→CA1 stimulation, respectively, and AAV1-Cre transsynaptic tracing was used to assess CA3→CA1 maturation. **(B and C)** High-frequency stimulation increased CA3→CA1 labeling of PV^-^ neurons (unpaired *t*-test, *t*_14_ = 2.58, *P* = 0.0220) and PV^+^ interneurons at P20 (unpaired *t*-test, *t*_14_ = 5.12, *P* = 0.0002). **(D and E)** Low-frequency stimulation reduced CA3→CA1 labeling of PV^-^ neurons (unpaired *t*-test, *t*_14_ = 2.26, *P* = 0.0403) and PV^+^ interneurons at P24 (unpaired *t*-test, *t*_14_ = 4.56, *P* = 0.0004). **(F)** Representative PNN and transsynaptic YFP labeling in CA1. **A**rrows indicate co-labeled cells. **(G and H)** High-frequency stimulation increased PNN assembly around PV^+^ interneurons at P20 (unpaired t-test, *t*_14_ = 6.22, *P* < 0.0001), with the increase preferentially occurring in transsynaptically labeled PV^+^ interneurons (ANOVA, projection × photostimulation: *F*_1,14_ = 19.31, *P* = 0.0006). **(I and J)** Low-frequency stimulation reduced PNN assembly around PV^+^ interneurons at P24 (unpaired *t*-test, *t*_14_ = 8.19, *P* < 0.0001), with the reduction preferentially occurring in transsynaptically labeled PV^+^ interneurons (ANOVA, projection × photostimulation: *F*_1,14_ = 5.05, *P* = 0.0413). **(K)** CA3→CA1 activity was increased or reduced during development before contextual fear conditioning at P20, P24, or P60. **(L)** High-frequency CA3→CA1 stimulation induced context-specific freezing at P20 but had no effect at P24 or P60; MEC→CA1 stimulation had no effect at any age (ANOVA, age × photostimulation: *F*_4,63_ = 5.95, *P* = 0.0004). **(M)** Low-frequency CA3→CA1 stimulation caused P24 mice to generalize across contexts but had no effect at P20 or P60; MEC→CA1 stimulation had no effect at any age (ANOVA, age × photostimulation: *F*_4,64_ = 4.65, *P* = 0.0023). **(N)** Experimental design for testing whether PNN maturation is required for the effects of increased TSP activity. ΔHAPLN1 was overexpressed in CA1 before high-frequency CA3→CA1 stimulation and contextual fear conditioning at P20. **(O and P)** High-frequency CA3→CA1 stimulation induced context-specific freezing at P20, which was prevented by ΔHAPLN1 (ANOVA, photostimulation × virus × context: *F*_1,28_ = 16.74, *P* = 0.0003), (ANOVA, photostimulation × virus: *F*_1,28_ = 29.26, *P* < 0.0001). **(Q)** High-frequency CA3→CA1 stimulation increased PNN assembly around CA1 PV^+^ interneurons, which was prevented by ΔHAPLN1 overexpression (ANOVA, photostimulation × virus: *F*_1,20_ = 19.30, *P* = 0.0003). **(R)** Experimental design for testing whether promoting PNN maturation can overcome reduced TSP activity. HAPLN1 was overexpressed in CA1 before low-frequency CA3→CA1 stimulation and contextual fear conditioning at P24. **(S and T)** Low-frequency CA3→CA1 stimulation caused mice to generalize across contexts at P24, whereas HAPLN1 overexpression restored context-specific freezing (ANOVA, photostimulation × virus × context: *F*_1,28_ = 13.29, *P* = 0.0011), (ANOVA, photostimulation × virus: *F*_1,28_ = 22.77, *P* < 0.0001). **(U)** Low-frequency CA3→CA1 stimulation reduced PNN assembly around CA1 PV^+^ interneurons, which was restored by wild-type HAPLN1 (ANOVA, photostimulation × virus: *F*_1,20_ = 18.51, *P* = 0.0003). Data are individual mice, mean ± s.e.m. Scale bars: white = 100 µm. \**P* < 0.05; ** *P* < 0.01; \*\*\**P* < 0.001; \*\*\*\**P* < 0.0001.

Given the developmental coupling between TSP innervation and PV interneuron maturation, we examined whether TSP activity also regulates PNN assembly around its downstream PV interneurons. High-frequency stimulation of CA3→CA1 terminals increased PNNs around CA1 PV^+^ neurons in P20 mice **(Fig. 5G, Fig. S6B, E).** Conversely, low-frequency stimulation of CA3→CA1 terminals decreased PNNs in P24 mice **(Fig. 5I, Fig. S6M, S)**. When combined with AAV1-Cre transsynaptic tracing, these changes were selective for transsynaptically labeled PV interneurons (**Fig. 5H, J**). Thus, activity within the developing TSP promotes PNN assembly selectively around synaptically connected PV interneurons, linking pathway activity to the maturation of its postsynaptic inhibitory targets.

We then examined the behavioral consequences of manipulating TSP or MSP activity during development. To increase activity, we delivered high-frequency stimulation of CA3→CA1 or MEC→CA1 terminals once daily for three consecutive days during different developmental windows (P17–19, P21–23, or P57–59), followed by contextual fear conditioning at P20, P24, or P60, respectively (**Fig. 5K**). Whereas control mice exhibited generalized memory, high-frequency stimulation of CA3→CA1 terminals, but not MEC→CA1 terminals, induced precocious context-specific freezing at P20, consistent with increased memory resolution (**Fig. 5L**, **Fig. S6C-D**). This effect was developmentally restricted, as identical stimulation had no effect on memory resolution at P24 or P60 (**Fig. 5L**, **Fig. S6F-G, I-J**).

Reducing TSP, but not MSP, activity produced the opposite developmental phenotype. CA3→CA1 terminals were stimulated with a low-frequency protocol for five consecutive days before contextual fear conditioning. Low-frequency stimulation of CA3→CA1 terminals had no effect at P20, when memory is normally generalized, but caused P24 mice to generalize across contexts, indicating delayed emergence of high-resolution memory (**Fig. 5M**, **Fig. S6Q-R**). This developmental delay was not permanent: mice receiving low-frequency stimulation of CA3→CA1 terminals from P19–P23 exhibited normal high-resolution memory when subsequently trained at P60 (**Fig. S7A-C**). Moreover, low-frequency stimulation in adulthood had no effect on memory resolution (**Fig. 5M**, **Fig. S6T-U**). Thus, TSP activity bidirectionally regulates pathway maturation, PNN assembly, and the developmental trajectory of memory resolution, with effects restricted to a sensitive period of hippocampal development.

Finally, we asked whether PNN maturation provides the causal link between developmental TSP activity and memory resolution. In our previous work, we developed viral tools targeting hyaluranon and proteoglycan link protein 1 (HAPLN1) that enable bidirectional manipulatation of PNN integrity: overexpression of wild-type HAPLN1 accelerates PNN assembly in developing mice, whereas overexpression of a dominant-negative HAPLN1 isoform (ΔHAPLN1) destabilizes PNNs (*11*) (**Fig. S8A-B**). Using these tools, we tested whether manipulating PNN maturational state could override the effects of experimentally accelerating or delaying TSP maturation. Overexpression of ΔHAPLN1 in CA1 prevented the precocious emergence of high-resolution memory at P20 induced by high-frequency CA3→CA1 terminal stimulation (**Fig. 5N-Q**). Conversely, overexpression of wild-type HAPLN1 restored high-resolution memory at P24, despite prior low-frequency CA3→CA1 terminal stimulation (**Fig. 5R-U**). These experiments establish PNN assembly as a critical downstream mechanism through which activity-dependent TSP maturation regulates the emergence of memory resolution.

## TSP activity mediates the effects of early-life experience on memory resolution

ELE and ELA affect many neural circuits (*38*, *41*). We therefore asked whether their effects on memory development are mediated specifically through activity-dependent maturation of the TSP. To address this question, we combined environmental manipulations with pathway-specific optogenetic stimulation and first examined maturation of CA3→CA1 inputs (**Fig. 6A**). In mice raised under ELE conditions, low-frequency stimulation of CA3→CA1 terminals prevented the enrichment-induced increases in both transsynaptic labeling of CA1 neurons, including PV^+^ neurons (**Fig. 6B-C**), and PNN assembly around CA1 PV^+^ neurons (**Fig. 6D**). Conversely, in mice that experienced ELA, high-frequency stimulation rescued the reduction in transsynaptic labeling associated with delayed TSP maturation (**Fig. 6E-F**) and restored PNNs around CA1 PV^+^ neurons (**Fig. 6G**). Thus, bidirectional manipulation of TSP activity was sufficient to override the effects of early-life experience on both pathway and PNN maturation.

**Fig. 6.**
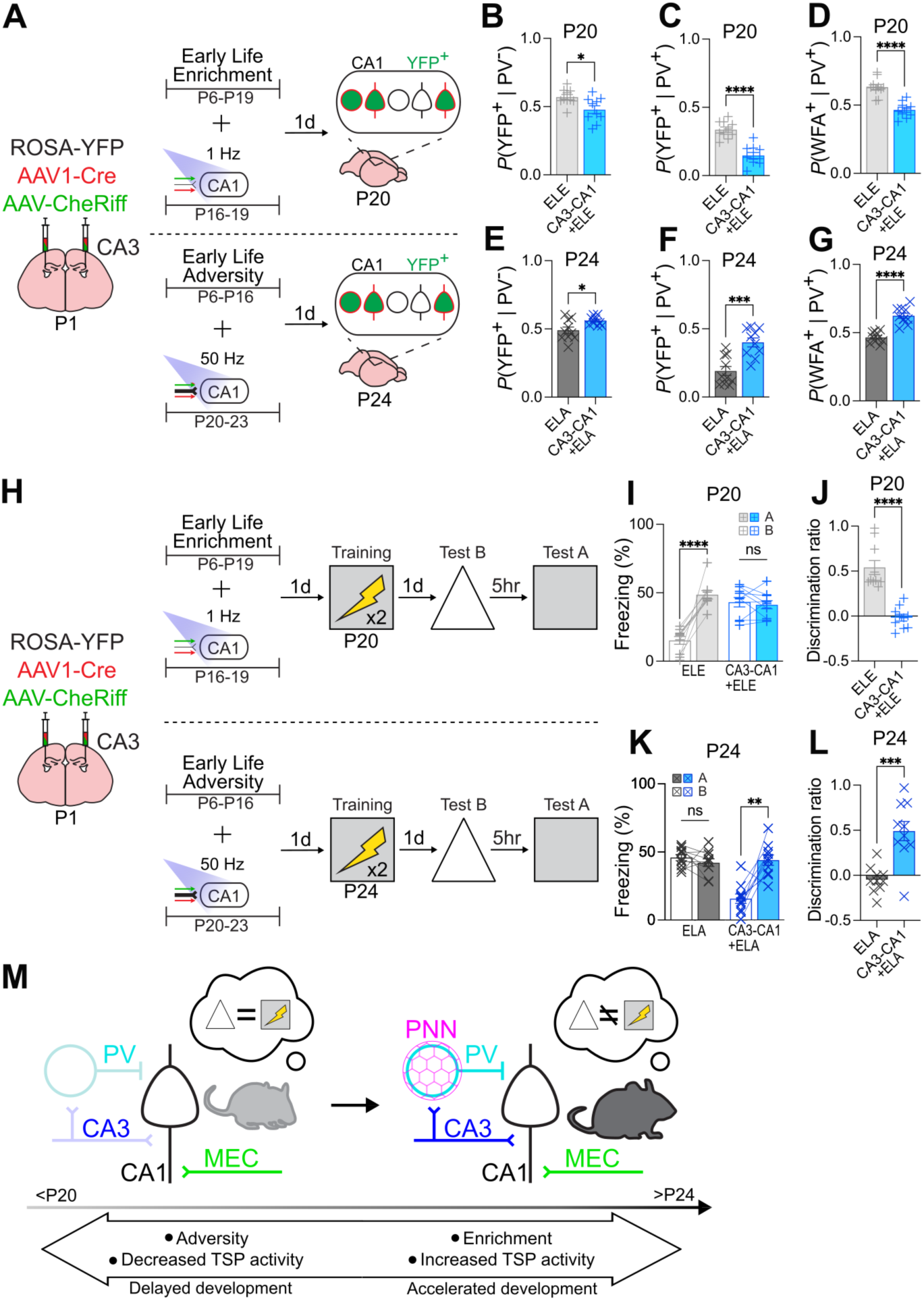
TSP activity mediates the effects of early-life experience on memory resolution. **(A)** ELE and ELA were combined with low-or high-frequency CA3→CA1 stimulation, respectively, and AAV1-Cre transsynaptic tracing to assess CA3→CA1 maturation. **(B to D)** Low-frequency stimulation prevented ELE-induced increases in CA3→CA1 labeling of PV**^-^** neurons (unpaired *t*-test, *t*_18_ = 2.70, *P* = 0.0148) and PV^+^ interneurons (*t*_18_ = 6.43, *P* < 0.0001), as well as increased PNN assembly around PV^+^ interneurons (*t*_14_ = 5.98, *P* < 0.0001). **(E to G)** High-frequency stimulation rescued ELA-induced reductions in CA3→CA1 labeling of PV**^-^** neurons (unpaired *t*-test, *t*_18_ = 2.70, *P* = 0.0145) and PV^+^ interneurons (*t*_18_ = 4.51, *P* = 0.0003), as well as reduced PNN assembly around PV^+^ interneurons (*t*_18_ = 6.87, *P* < 0.0001). **(H)** ELE and ELA were combined with low- or high-frequency CA3→CA1 stimulation, respectively, before contextual fear conditioning. **(I and J)** P20 ELE mice showed context-specific freezing, which was prevented by low-frequency CA3→CA1 stimulation (ANOVA, stimulation × context: *F*_1,18_ = 38.57, *P* < 0.0001), reducing context discrimination (unpaired *t*-test, *t*_18_ = 6.45, *P* < 0.0001). **(K and L)** P24 ELA mice generalized across contexts, whereas high-frequency CA3→CA1 stimulation restored context-specific freezing (ANOVA, stimulation × context: *F*_1,18_ = 18.66, *P* = 0.0004) and context discrimination (unpaired *t*-test, *t*_18_ = 4.63, *P* = 0.0002). **(M)** Model in which early-life experience regulates TSP activity and maturation, PNN assembly around CA1 PV interneurons, and the emergence of high-resolution memory. Data are individual mice, mean ± s.e.m. \**P* < 0.05; ** *P* < 0.01; \*\*\**P* < 0.001; \*\*\*\**P* < 0.0001.

We then determined whether these manipulations also reversed the effects of early-life experience on memory resolution (**Fig. 6H**). As expected, ELE caused P20 mice to freeze selectively in the training context, an effect abolished by low-frequency stimulation that delayed CA3→CA1 maturation (**Fig. 6I–J**). In contrast, ELA caused P24 mice to generalize across contexts, an effect reversed by high-frequency stimulation that restored age-appropriate CA3→CA1 maturation (**Fig. 6K–L**). Together, these findings indicate that the effects of early-life experience on the developmental trajectory of memory resolution are mediated through activity-dependent regulation of TSP maturation.

## Discussion

Here, we identify a circuit mechanism underlying the developmental emergence of high-resolution memory. Consistent with previous studies, we found that the ability to form high-resolution memories emerges during the fourth postnatal week in mice. Before this transition, the relatively mature MSP supports the formation of low-resolution memories, confirming previous predictions (*10*, *28*, *42*). In contrast, the TSP matures later, with a particularly pronounced developmental increase in CA3→CA1 innervation of PV interneurons between P20 and P24. This maturation was accompanied by the functional recruitment of CA1 PV interneurons during learning and was necessary for the emergence of high-resolution memory. Manipulating TSP activity or early-life experience bidirectionally shifted pathway maturation, PNN assembly, PV recruitment, and the developmental emergence of high-resolution memory. Together, these findings identify the sequential maturation of hippocampal input pathways, and particularly the recruitment of CA1 inhibitory circuitry by the developing TSP, as a key mechanism underlying the transition from generalized to context-specific memory (**Fig. 6M**).

These findings reconcile two previously distinct accounts of the development of high-resolution memory. Our previous work focused on local maturational events within CA1, identifying a cellular mechanism for the developmental transition from generalized to high-resolution memory. Between P20 and P24, PNNs assemble around CA1 PV interneurons, promoting their functional maturation and recruitment during learning. Increased PV interneuron-mediated inhibition, in turn, constrains the size of CA1 encoding ensembles, producing the sparse representations associated with high-resolution memory. Importantly, manipulating PNN maturation bidirectionally shifts PV interneuron recruitment, ensemble sparsity, and memory resolution, establishing a causal relationship between these developmental events occurring locally in CA1 (*11*). This cellular account has remained largely separate from a prominent circuit-level account of memory development, which proposes that generalized and high-resolution memories reflect the differential maturation of the MSP and TSP, respectively (*10*, *42*, *43*). The present findings bridge these levels of explanation. We show that maturation of the TSP promotes PNN assembly and consequently increases recruitment of CA1 PV interneurons. The relationship between TSP and PNN maturation in CA1 is likely reciprocal—while developmental CA3→CA1 activity is required for PNN assembly, as shown here, maintenance of mature CA3→CA1 inputs to PV interneurons likely depends on lecticans found in the PNN for their stabilization (e.g., brevican) (*44*, *45*). Taken together, our findings link the late-developing hippocampal pathway proposed by cognitive neuroscience models to the cellular mechanisms that drive the emergence of sparse, high-resolution memory.

More broadly, our findings suggest that the emergence of high-resolution memory reflects a sensitive period in hippocampal circuit development. Canonical sensitive-period mechanisms have been most extensively characterized in developing sensory cortices, including visual and somatosensory cortex, where experience shapes circuit organization through activity-dependent maturation of inhibitory networks (*46*). In particular, maturation of PV interneurons by the assembly of extracellular PNNs is a central regulator of the timing and expression of these periods of heightened developmental plasticity (*12*, *31*, *47*). Similar mechanisms have been proposed to operate in later-developing brain regions supporting cognitive functions, including learning and memory, raising the possibility that principles first identified in sensory systems are conserved across cortical and hippocampal development (*48–50*). The developmental transition identified here exhibits several of these defining features. Early-life enrichment and adversity bidirectionally shifted TSP maturation, PNN assembly and PV interneuron recruitment, and the onset of high-resolution memory, while directly manipulating TSP activity was sufficient to reproduce or reverse these experience-dependent effects. Notably, this plasticity was developmentally restricted: increasing TSP activity accelerated the emergence of high-resolution memory in juvenile mice, whereas reducing TSP activity delayed this transition, but comparable manipulations were ineffective once the circuit had reached maturity. Thus, the TSP may play two interrelated but distinct roles across the lifespan. In the mature hippocampus, it provides a circuit substrate for forming high-resolution representations; during development, activity within this still-maturing pathway appears to instruct maturation of the CA1 inhibitory circuitry that ultimately enables these computations.

The sensitivity of TSP maturation to early-life experience provides a potential circuit mechanism through which early-life environments shape the development of memory. We previously found that early-life adversity delays, whereas enrichment accelerates, the emergence of high-resolution memory by altering the pace of PNN maturation in CA1 (*31*). Here, we identify activity within the developing TSP as an instructive signal for this maturational program. Bidirectional manipulation of TSP activity shifted both CA3→CA1 innervation of PV interneurons and PNN maturation, linking activity in this pathway to the development of local inhibitory circuitry. Critically, manipulating TSP activity was sufficient to override the effects of enrichment and adversity on both circuit maturation and memory precision, indicating that these early-life experiences regulate memory development through their effects on the developing TSP.

Together, our findings identify activity-dependent maturation of the TSP as a key developmental process linking early-life experience to the emergence of high-resolution memory. More broadly, they suggest that the timing of hippocampal maturation is itself experience-dependent, providing a mechanism through which early environments may shape memory development. This possibility is particularly relevant to human development, where socioeconomic disadvantage and early-life adversity are associated with poorer episodic memory, whereas enriched early experiences, including high-quality parental care and early education, are associated with better memory outcomes (*51*, *52*). Our findings raise the possibility that such bidirectional effects of early-life experience may arise, in part, by shifting the developmental trajectory of hippocampal circuits that support high-resolution memory.

## Acknowledgments

We thank Antonietta Decristofaro, Daisy Lin, Mika Yamamoto for their technical assistance. We also thank members of the Frankland and Josselyn labs for general comments.

## Funding

Canadian Institutes of Health Research (PJT-180538)

Ontario Graduate Scholarship (C.M.)

SickKids Restracomp Fellowship (C.M., A.I.R.)

Canadian Neurodevelopmental Research 15 Training Platform (A.I.R.)

## Author contributions

Conceptualization: C.M., A.I.R., P.W.F.

Experimentation: C.M., M.S., S.H.

Statistical analysis: C.M.

Writing: C.M., A.I.R., P.W.F.

Feedback and editing: S.A.J.

## Competing interests

The authors declare no competing interests.

## Data and materials availability

Data are available from the corresponding author on request.

## Supplementary Materials

### Materials and Methods

#### Mice

Three mouse lines were used. Male and female F1 hybrid (C57BL/6NTac x 129S6/SvEvTac) wild-type (WT) mice were used for all experiments where not otherwise stated. Rosa-YFP mice (B6;pNestinCRE x ROSA26YFP) which express the YFP protein in the presence of Cre recombinase were used for anterograde tracing (*53*). Rosa-YFP breeders were maintained on a C57BL/6 background and crossed with 129S6/SvEvTac breeders to generate the F1 hybrid Rosa-YFP mice used in experiments. PV-Cre knock-in driver transgenic mice (B6;129P2-Pvalbtm1(cre)Arbr/J, Strain# 008069) which express Cre recombinase in PV⁺ interneurons, without disrupting endogenous PV expression, were originally generated by Silvia Arber, FMI, and obtained from Jackson Laboratory (*54*). These mice were used to test the effect of chemogenetically exciting PV^+^ interneurons in combination with inhibiting CA3 to CA1 projections. Homozygous PV-Cre mouse breeders were maintained on a C57BL/6 background and crossed with 129S6/SvEvTac breeders to generate the F1 hybrid PV-Cre mice used in experiments.

Mice were bred at the Hospital for Sick Children and group-housed on a 12-h light/dark cycle with food and water ad libitum. All experiments took place during the light phase. Litter birthdate was designated postnatal day (P) 0. Litters ranged in size from 5-12 pups. Except where noted (see *Experiential Interventions* section), experimental mice were housed in breeding cages (identical to standard housing cages, 19 cm x 28 cm x 15 cm) with the dam from P0-20 and then weaned on P21 with same-sex littermates, with 2-5 mice per cage. Except where noted (see *Experiential Interventions*), all cages contained 80 g of cob bedding and 8 g of nesting material, and breeding cages contained one hut or dome. Mice used for behavioral experiments on P20-21 remained in the breeding cage with the dam for the duration of the study. Each experimental condition contained mice derived from 2-6 separate litters with a maximum of 2 males and 2 females from each litter in a single experimental condition. All procedures were approved by the Hospital for Sick Children Animal Care and Use Committee and conducted in accordance with Canadian Council on Animal Care.

#### Drugs

*DREADD agonist 21 (C21)*. C21 dihydrochloride (Tocris, cat# 6422) was prepared as a stock solution of 10 mg/ml in dH2O and stored at -20 °C. Stock solution was later thawed and diluted 1:100 in PBS immediately before use. Diluted C21 was administered via i.p. injection 1 h before contextual fear training (1.0 mg/kg) to activate DREADD-expressing neurons.

#### Viruses

##### HSV

Replication-defective Herpes Simplex Viruses (HSV) were used to transiently express transgenes in hippocampal neurons. HSV is naturally neurotrophic and enables transgene expression that peaks and dissipates 3-4 d and 10 d after microinjection, respectively, as shown previously in CA1 of developing mice (*11*, *31*). Expression of transgenes was driven by the endogenous HSV promoter IE4/5 (except for HSV-GFP which used the CMV promoter). HSV virus was packaged in-house using replication-defective helper virus, purified on a sucrose gradient, pelleted, and resuspended in 10% sucrose. Final titers of HSVs were approximately 10^6^ infectious units/ml. The following HSV constructs were used:

##### HSV-NpACY

We used HSV-NpACY to bidirectionally manipulate the activity of sparse subsets of neurons. HSV-NpACY contains both enhanced channelrhodopsin-2 (ChR2-H134R) fused to enhanced yellow fluorescent protein (eYFP) and halorhodopsin 3.0 (NpHR3.0). These opsins are spectrally compatible, allowing for neuronal excitation by ChR2 with blue light (473 nm) and neuronal silencing by NpHR3.0 with red light (660 nm) (*55*).Opsin genes were connected in the viral vector using a 2A self-cleavage linker (p2A).

##### HSV-ΔHapln1

We used HSV-ΔHapln1 to manipulate PNNs. HSV-ΔHapln1 expresses a dominant-negative HAPLN1 link protein (ΔHAPLN1) fused with cysteine-free GFP. ΔHAPLN1 maintains the wild-type C-terminus domain that binds hyaluranon in the extracellular matrix but lacks the N-terminus domain that binds CSPGs. We previously generated the *ΔHapln1* construct (Addgene #200778; pAAV-ΔHapln1) and showed that viral expression of ΔHAPLN1 interferes with endogenous HAPLN1 function to destabilize PNNs locally in the brain (*11*). The relevant cDNA was removed from the AAV vector by restriction digest and cloned into the multiple cloning site of pHSV.

##### HSV-Hapln1

We used HSV-Hapln1 to manipulate PNNs. HSV-Hapln1 expresses the wild-type mouse HAPLN1 protein fused with cysteine-free GFP. We previously generated the *Hapln1* construct (Addgene #200777; pAAV-Hapln1) and showed that viral expression of HAPLN1 accelerates PNN formation in brains of juvenile mice (*11*). The relevant cDNA was removed from the AAV vector by restriction digest and cloned into the multiple cloning site of pHSV.

##### HSV-GFP

HSV-GFP (HSV-p1005) expresses the GFP fluorophore (*56*) and was used as a control virus.

##### AAV

We used adeno-associated viruses (AAV) to manipulate neuronal activity in a circuit specific manner. Transgene expression peaks 3-4 weeks after AAV microinjection and is relatively stable in the following weeks. Except for AAV1-Cre, which was purchased commercially, AAVs (DJ serotype) were generated in-house in HEK293T cells with the AAV-DJ Helper Free Packaging System (Cell Biolabs, Inc., cat# VPK-400-DJ) using the manufacturer-suggested protocol. Viral particles were purified using Virabind AAV Purification Kit (Cell Biolabs, Inc., cat# VPK-140). AAV titers were approximately 1.0 ×10^11^ infectious units/ml. The following AAV constructs were used:

##### AAV1-hSyn-Cre-WPRE-hGH (AAV1-Cre)

AAV1-Cre was a gift from James M. Wilson (Addgene plasmid # 10555-AAV1; http://n2t.net/addgene:105553; RRID:Addgene_105553). AAV1-Cre expresses large quantities of Cre recombinase in the synaptic terminal of infected neurons, some of which is transported anterogradely to post-synaptic neurons (*57*, *58*). We injected AAV-1 into CA3 or MEC of Rosa-YFP mice to label neurons in CA1 receiving synaptic input from CA3 or MEC, respectively. Expression was driven by the human synapsin (hSyn) promoter.

##### AAV(DJ)-CaMK2α-NpACY (AAV-NpACY)

We used AAV-NpACY to inhibit CA3 neurons projecting to CA1 in PV-Cre mice or to inhibit individual hippocampal subregions in WT mice. NpACY expressed by this construct was identical to that described for HSV-NpACY above. Expression of NpACY in principal neurons was driven by the CaMK2α promoter.

##### AAV(DJ)-CMV-GFP (AAV-GFP)

We used AAV-GFP as a control virus for AAV-NpACY. pAAV-CMV-GFP was a gift from Dr. Connie Cepko (Addgene plasmid # 67634; http://n2t.net/addgene:67634; RRID:Addgene_67634). Expression of GFP was driven by the CMV promoter.

##### AAV(DJ)-hSyn-DIO-hM3D(Gq)-mCherry (AAV-DIO-hM3Dq)

We used AAV-DIO-hM3Dq to excite PV-positive interneurons in the CA1 of PV-Cre mice. AAV-DIO-hM3Dq was a gift from Bryan Roth (Addgene plasmid # 44361 ; http://n2t.net/addgene:44361 ; RRID:Addgene_44361). Expression of hM3Dq-mCherry in Cre^+^ neurons was driven by the hSyn promoter.

##### AAV(DJ)-hsyn-CheRiff-eGFP (AAV-CheRiff)

We used AAV-CheRiff to manipulate activity of CA3 or MEC inputs to CA1. AAV-CheRiff was a gift from Adam Cohen (Addgene plasmid # 51697 ; http://n2t.net/addgene:51697 ; RRID:Addgene_51697). CheRiff is a blue light sensitive excitatory opsin with fast kinetics that enables light-dependent driving of neuronal firing up to 50 Hz (*59*).Expression CheRiff-eGFP was driven by the hSyn promoter.

#### Tracers

*Cholera Toxin subunit-b (CTb).* CTb is a retrograde tracer derived from the non-toxic subunit-b of cholera toxin. CTb conjugated to fluorophores was obtained from Invitrogen (CTb-488: cat# C34775; CTb-647: cat# C34778). CTb was dissolved in 0.04 M PBS to a concentration of 40 kM and stored in the dark at 4 °C until use.

### Surgery

Surgeries were performed at various ages. Except for neonatal surgeries (see below), mice were pre-treated with atropine sulfate (0.1 mg/kg, i.p.), anesthetized with chloral hydrate (400 mg/kg, i.p.), administered meloxicam (4 mg/kg, s.c.) for analgesia, and placed into stereotaxic frames. The scalp was incised and retracted, and holes were drilled above the target area. Unless otherwise specified, viruses were injected bilaterally via a glass micropipette connected via polyethylene tubing to a 15 *μ*l microsyringe (Hamilton) at a rate of 0.1 *μ*l/min and remained in place for an additional 5 min to ensure virus diffusion. Following microinjections, the scalp was sutured and polysporin was applied. Mice were administered 0.9% saline (0.5-1.0 ml, s.c.) and placed in a clean cage on a heating pad to recover. Recovering mice monitored until ambulatory, at which point they were returned to the colony. Animals were monitored twice daily for at least 3 days, and additional analgesic were delivered if needed.

For all behavioural experiments involving virus microinjections, only mice showing strong bilateral expression (i.e., expression limited to the target brain region and observable in a minimum of 3 brain sections) were included in the final data set. Additionally, for optogenetics experiments, only mice with optic fibers placed correctly above the region of interest were included in the final data set.

#### Neonatal virus microinjection

Neonatal virus injection procedures were adapted from previous studies (*11*, *31*). P1 mice were anesthetized by hypothermia and mounted to a chilled metal plate using laboratory tape. The scalp was incised and retracted, and the glass micropipette connected to a nanoliter injector (Nanoject III, Drummond Scientific) was used to pierce the skull above CA3 (approximate coordinates AP -0.7 mm, ML ±1.65 mm from bregma, DV -1.8 mm from skull surface), CA1 (AP +1.0 mm, ML ± 0.8 from lambda, DV -1.1 mm from skull surface), or MEC (AP-0.3 mm from lambda, ML ±2.0 mm from lambda, DV -1.5 mm from skull surface). 0.2 *μ*l AAV was injected over 1 min at the specified coordinates and remained in place for an additional 2 min to prevent reflux. The scalp was sealed with Vetbond Tissue Adhesive (3M) and covered with polysporin. For some experiments, paws were tattooed using non-toxic black ink for later mouse identification. The entire procedure was performed within 10-12 min. Pups were placed on a heating pad and continuously monitored until mobile. Pups remained on the heating pad until all surgeries were completed, at which time the entire litter was returned to the dam.

#### Other virus microinjection

The following coordinates and virus volumes were used for dorsal CA1: ≥P40, coordinates AP -1.8 mm, ML ±1.5 mm, DV -1.5 mm from bregma; P17-P21, coordinates AP -1.8 mm, ML ±1.45 mm, DV -1.5 mm from bregma. For dorsal DG: ≥P40, coordinates AP -2.5 mm, ML ±3.0 mm, DV -2.8 mm from bregma; P17-P21, coordinates AP -2.0 mm, ML ±1.5 mm, DV -2.1 mm from bregma. For dorsal CA3: ≥P40: coordinates AP -2.15 mm, ML ± 2.45 mm, DV -2.3 mm from bregma; For P17-21, coordinates AP -2.15 mm, ML ± 2.3 mm, DV -2.2 mm from bregma. For MEC: ≥P40, coordinates AP +0.2 mm, ML ±3.5 mm, DV -2.2 mm from transverse sinus, lambda, and skull surface, respectively; P17-P21, coordinates AP +0.2 mm, ML ±3.3 mm, DV -2.1 mm from transverse sinus, lambda, and skull surface, respectively.

#### Fiber implantation

Optical fibers were constructed by inserting and securing a length of optical fiber (200 *μ*m diameter, 0.37 NA) into a 1.25 mm zirconia ferrule such that the fiber extended 3 mm beyond the ferrule. These fibers were implanted 0.1 mm dorsal to the CA1, CA3, or MEC and secured to the skull with dental cement and 2-3 jeweler screws. Mice were allowed to recover for at least 48 hours before the experiment.

### Experiential Interventions

Two experiential interventions were used, referred to hereafter as early-life adversity (ELA) and early-life enrichment (ELE).

The ELA protocol consisted of limited bedding and nesting from P6-16 (inclusive). Homecages were reduced to one-quarter of typical resources (i.e., 20 g of cob bedding and 2 g of nesting material), and cages were changed every 3 days. ELA mice were housed in the resource-restricted cages until P16 when they were returned to standard housing conditions.

The ELE protocol was adapted from previous studies using mice. For the ELE intervention, all pups from the litter and the dam were relocated to a larger polycarbonate rat cage (enrichment cage, 25 cm x 40 cm x 20 cm) beginning on P6. The enrichment cages contained 3 huts and/or domes and 3-4 pieces of commercially purchased hamster tunnels, and twice the amount of nesting materials compared with controls (i.e., 16 g). Enrichment was rearranged or changed every 3 days, and enrichment cages were changed once per week. ELE mice were housed in the enrichment cages until P19 when they were returned to standard housing conditions. Control mice remained in standard housing cages during the equivalent period.

A subset of ELA, ELE, and control mice were weighed on P6, P13, P20, P24, and P60 and observed for eye opening on P11-14. Offspring survival was not affected by the experiential interventions, and there were no observable changes in mouse health in any of the conditions, other than modest changes in weight gain in ELE and ELA mice (Fig. S4).

### Behaviour

#### Fear conditioning

Contextual fear conditioning occurred in test chambers (31 × 24 × 21 cm) with shock-grid floors (Med Associates). During training, mice were placed in the chambers for 2 min, after which two foot shocks (0.5 mA, 2 s duration, 1 min apart) were delivered. Mice were removed from the chambers 1 min after the last shock (4 min total). The next day, mice were placed in a similar but novel chamber (Context B) for a 5 min test. 5 h later the mice were placed in the original training chamber (Context A) for another 5 min test. Context B was approximately the same size as Context A, with white plastic floor and triangular white plastic walls. Context C was a rectangular chamber (15 × 45 × 25 cm) with an open top, made of white plexiglass, and it was located in a separate room. Mouse behavior was recorded with overhead cameras and FreezeFrame v.3.32 software (Actimetrics). For contextual fear memory tests, the amount of time mice spent freezing was scored during the entire test session automatically using FreezeFrame software or manually by a researcher blinded to the conditions (when optic patch cables were attached). Freezing was defined as the cessation of movement, except for breathing (*60*).

#### Optogenetic stimulation

Mice used in the optogenetics experiments were habituated to the optic patch cables for 5 min one day before before training. For optogenetic inhibition during training mice were attached to the patch cables and placed in Context A. For the duration of the training period as described above, continuous red (660 nm, 7-10 mW) light was delivered. For optogenetic stimulation to induce c-Fos expression mice were habituated to the experimental room for 1 hr before being attached to the optic patch cables and placed into a clean cage. Mice received blue light stimulation (473 nm, 1 mW, 20 Hz, 15-ms pulses) for 90 s.

#### High and low frequency stimulation

For high frequency stimulation, mice were placed alone in their homecage and received 5 trains of blue light stimulation (473 nm, 10 mW, 50 Hz, 2 ms per pulse, 75 pulses/train) separated by a 3 min interval. For low frequency stimulation, mice received weak paired-pulse stimulations at an interval of 1 Hz (473 nm, 1 mW, 1 Hz, 2 ms per pulse, 900 paired pulses at 200 ms intervals). Each session lasted approximately 15 min. Mice received one stimulation session per day for 3 (high frequency stimulation) or 5 days (low frequency stimulation).

### Histology

#### Perfusion and tissue preparation

At the appropriate age, following the appropriate delay after behavior experiments, mice were transcardially perfused with chilled 1× PBS followed by 4% paraformaldehyde (PFA), fixed in PFA overnight at 4 °C, and transferred to 30% sucrose solution for at least 72 h. When appropriate, perfusions occurred 90 min after training. Brains were sectioned coronally or sagittally using a cryostat (Leica CM1850), and a 1⁄4 sampling fraction was used to obtain 4 sets of 50-*μ*m sections. Sections for immunohistochemistry were stored at 4 °C in PBS containing 0.1% NaN_3_ until staining.

#### Immunohistochemisty

Immunofluorescence staining was conducted as previously described (*11*, *61*). For experiments involving quantification of the number of cells positive for immunofluorescence, all staining was performed at once using the same antibody solutions. Free-floating sections were blocked with PBS containing 4% normal goat serum and 0.5% Triton-X for 1 h at room temperature. Afterwards, sections were incubated with primary antibodies in fresh blocking solution: rabbit anti-c-Fos (1:500, Synaptic Systems, cat#226003), rat anti-c-Fos (Synaptic Systems, cat #226017), chicken anti-GFP (1:1000, Aves, cat# GFP-1010), rabbit anti-RFP (1:1000, Rockland Immunochemicals, cat# 600-401-379), mouse anti-PV (1:1000, Sigma, cat# 1572), rabbit anti-SST (1:250, Synaptic Systems, cat#257003), biotinylated WFA (1:1000, Sigma, cat# L1516) for 24 h or 48 h (for c-Fos and PV experiments) at 4 °C. Sections were washed three times for 15 min each with PBS containing 0.1% Tween-20 (PBST), then incubated with PBST containing secondary antibodies: goat anti-chicken Alexa Fluor 488 (1:500, Invitrogen, cat# A-11039), goat anti-mouse Alexa Fluor 647 (1:500, Invitrogen, cat# A-21235), goat anti-rabbit Alexa Fluor 568 (1:500, Invitrogen, cat# A-11011), goat anti-rat Alexa Fluor 633 (1:500, Invitrogen, cat# A-21094), streptavidin Alexa Fluor 568 (1:500, Invitrogen, cat# S-11226) for 2 h at room temperature. Sections were washed with PBS, counterstained with DAPI, mounted on gel-coated slides, and coverslipped with Permafluor mounting medium (ThermoFisher Scientific, cat# TA-030-FM).

#### Imaging

Images were obtained using a confocal laser scanning microscope (LSM 710; Zeiss). For visualization of virus expression, images were acquired with a 20× objective. For image quantification, z-stacks were acquired using a 40× objective (N.A. = 1.3; 15-40 slices with a 1 *μ*m step size). For all experiments involving quantification of the number of cells positive for immunofluorescence, all images were acquired using identical imaging parameters (laser power, photomultiplier gain, pinhole, and detection filter settings) in a minimal number of imaging sessions (when possible, in one session). For each experiment, imaging parameters were set using a sample section from a control mouse.

#### Quantification

For cell counting experiments in the CA1 every fourth section was assessed for the marker(s) of interest. For each mouse, cells were counted manually in Fiji (National Institutes of Health) using 5 images acquired from 3-5 sections and averaged. To estimate the number of DAPI+ cells in the CA1 pyramidal layer, DAPI^+^ cells were counted in a small volume (approximately 12-20 ×10^3^ *μ*m^3^) to obtain the DAPI^+^ density (mm^3^). For all experiments, the volume of the pyramidal layer within each image was measured and multiplied by the DAPI^+^ density value for the appropriate age group to obtain the estimated DAPI^+^ cell number. Overlap between different markers are reported as conditional probabilities. The proportion of PV***^-^*** pyramidal layer cells (putative excitatory population) expressing c-Fos or YFP are reported as *P*(c-Fos^+^|PV^-^) or *P*(YFP^+^| PV^-^), and the proportion of PV***^+^*** cells expressing c-Fos or YFP are reported as P(c-Fos^+^|PV^+^) or *P*(YFP^+^|PV^+^). Colocalization of the PNN marker WFA with PV-positive interneurons is reported as *P*(WFA^+^|PV^+^).

### Statistical analysis

No statistical tests were used to predetermine sample sizes, but our sample sizes were similar to those reported in previous publications (*11*, *61*). Mice were pseudo-randomly assigned to all groups to achieve roughly equivalent group sizes. During data collection and quantification, experimenters were blinded to group assignment, except for age and testing context (during data collection) and testing context (during quantification), which could not be blinded.

Data were analyzed using unpaired *t*-tests or analysis of variance (ANOVA) with repeated measures when appropriate. When appropriate, ANOVAs were followed by Tukey’s post-hoc comparisons. In some experiments (Fig. 2,3,4,5), control mice were pooled into a single control group. Sub-group means were not statistically different from one another (data not shown). Both male and female mice were used in all experiments, and no differences were found between the two groups unless otherwise reported. Statistical significance was set at *P* < 0.05. Statistical analyses were performed in Graphpad Prism (version 10.0.1) or using customs scripts in R.

## Supplementary Figures

**Fig. S1.**
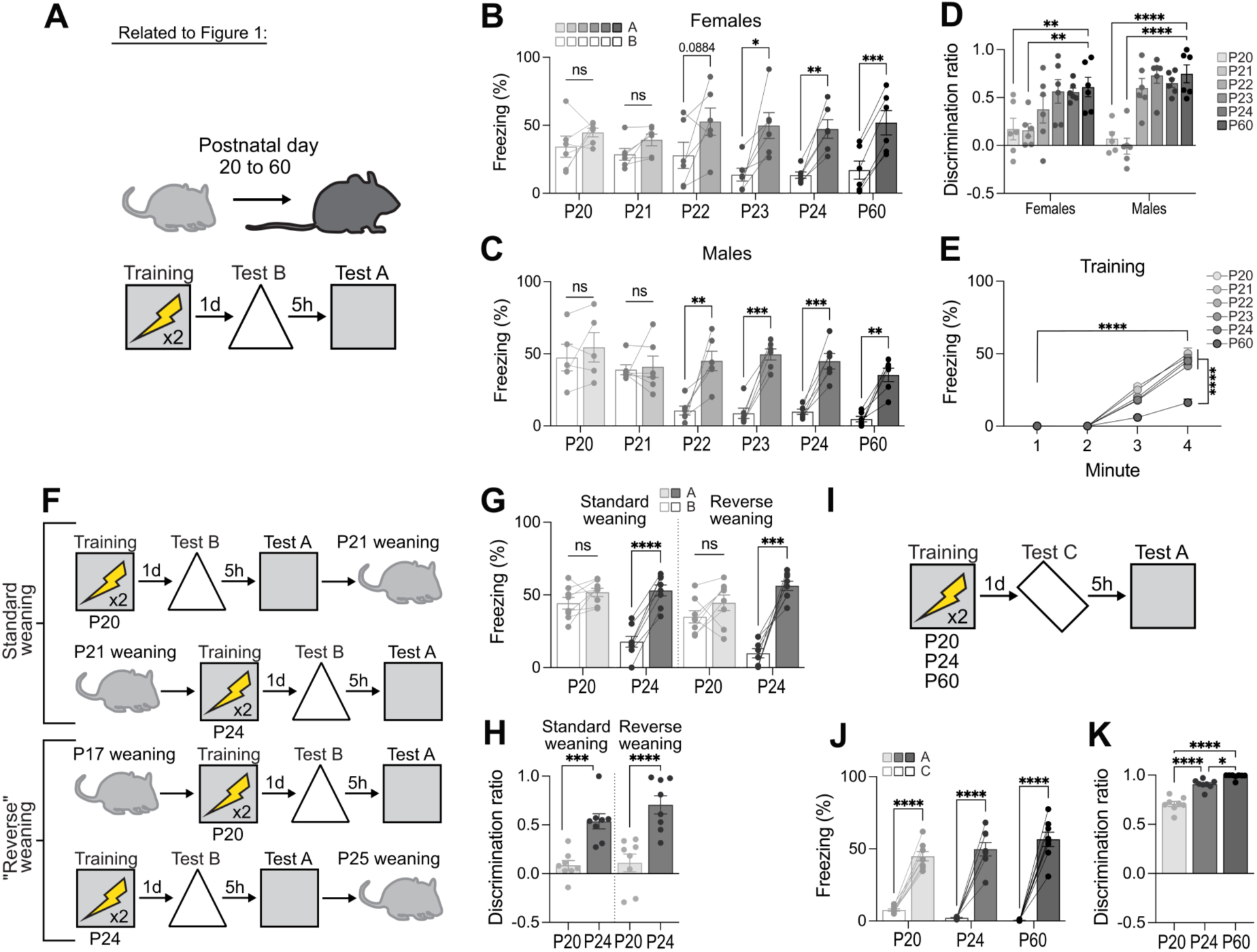
Developmental emergence of high-resolution memory is independent of sex, weaning, and perceptual ability (Related to Figure 1). **(A)** Experimental design for assessing the developmental emergence of high-resolution contextual memory. **(B, C, and D)** The transition from contextual generalization to context specific freezing occurred between P21 and P22 in both females (ANOVA, age × context: *F*_5,30_ = 7.38, *P* < 0.0001) and males (ANOVA, age × context: *F*_5,29_ = 6.23, *P* = 0.0005) which is also reflected by discrimination ratios (ANOVA, age: *F*_5,59_ = 16.29, *P* < 0.0001; age × sex: *F*_5,59_ = 1.37, *P* = 0.2476). **(E)** Mice at all ages show learning during contextual fear conditioning, although P60 mice freeze less than P20-P24 mice (ANOVA, age × minute: *F*_15,195_ = 14.98, *P* < 0.0001). **(F)** Reversed and standard weaning protocols for P20 and P24 mice. **(G and H)** Freezing (ANOVA, age × context × weaning: *F*_1,28_ = 0.74, *P* = 0.7353) and discrimination (ANOVA, age × weaning: *F*_1,28_ = 0.81, *P* = 0.3754) in P20 and P24 mice was not affected by weaning protocol. **(I)** Contextual fear generalization test with a highly distinct context C. **(J and K)** Mice at all ages showed context specific freezing (ANOVA, age × context: *F*_2,21_ = 4.79, *P* = 0.0194) and high discrimination (ANOVA, age: *F*_2,21_ = 61.15, *P* < 0.0001) when tested in context C. Data points are individual mice with mean ± s.e.m. \**P* < 0.05; ** *P* < 0.01; \*\*\**P* < 0.001; \*\*\*\**P* < 0.0001.

**Fig. S2.**
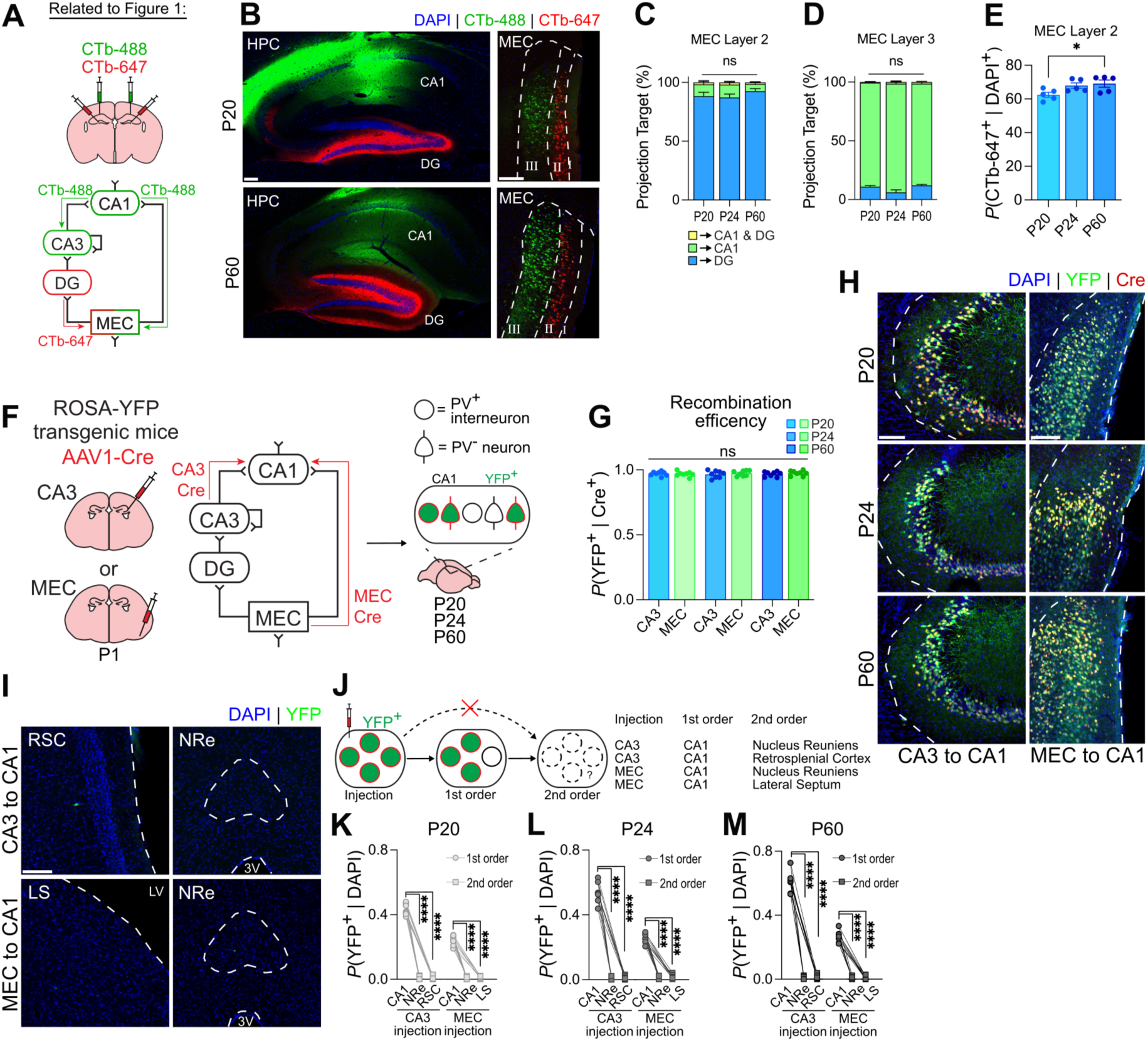
Validation of pathway-specific retrograde and anterograde transsynaptic tracing (Related to Figure 1). **(A and B)** Retrograde tracing from CA1 and dentate gyrus (DG) with CTb-488 and CTb-647 at P20, P24, and P60. **(C and D)** Distinct layer-specific projections from the MEC layer II (ANOVA, age × projection: *F*_4,24_ = 0.94, *P* = 0.4605) and layer III (ANOVA, age × projection: *F*_4,24_ = 4.07, *P* = 0.0117). (E) Projections from MEC layer II to DG are reduced at P20 compared to P60 (ANOVA, age: *F*_2,12_ = 4.14, *P* = 0.0429). **(F)** Validation of AAV1-Cre transsynaptic tracing in Cre-dependent ROSA-YFP reporter mice. AAV1-Cre was injected into CA3 or MEC, and YFP labeling was quantified at the injection site. **(G)** YFP labeling in Cre-positive neurons was observed following injections into both regions across development (ANOVA, age × region: *F*_2,42_ = 0.33, *P* = 0.7221). **(H)** Representative image of YFP expression in Cre-positive neurons in CA3 and MEC. **(I to M)** AAV1-Cre injections in CA3 and MEC resulted in robust expression in the monosynaptically connected CA1 but not in disynaptic (i.e., second order) downstream targets at P20 (ANOVA, projection: *F*_1,17_ = 1263.0, *P* < 0.0001), P24 (ANOVA, projection: *F*_1,16_ = 860.6, *P* < 0.0001), or P60 (ANOVA, projection: *F*_1,15_ = 983.0, *P* < 0.0001). Data points are individual mice with mean ± s.e.m. Scale bars: white = 100 µm. \**P* < 0.05; ** *P* < 0.01; \*\*\**P* < 0.001; \*\*\*\**P* < 0.0001.

**Fig. S3.**
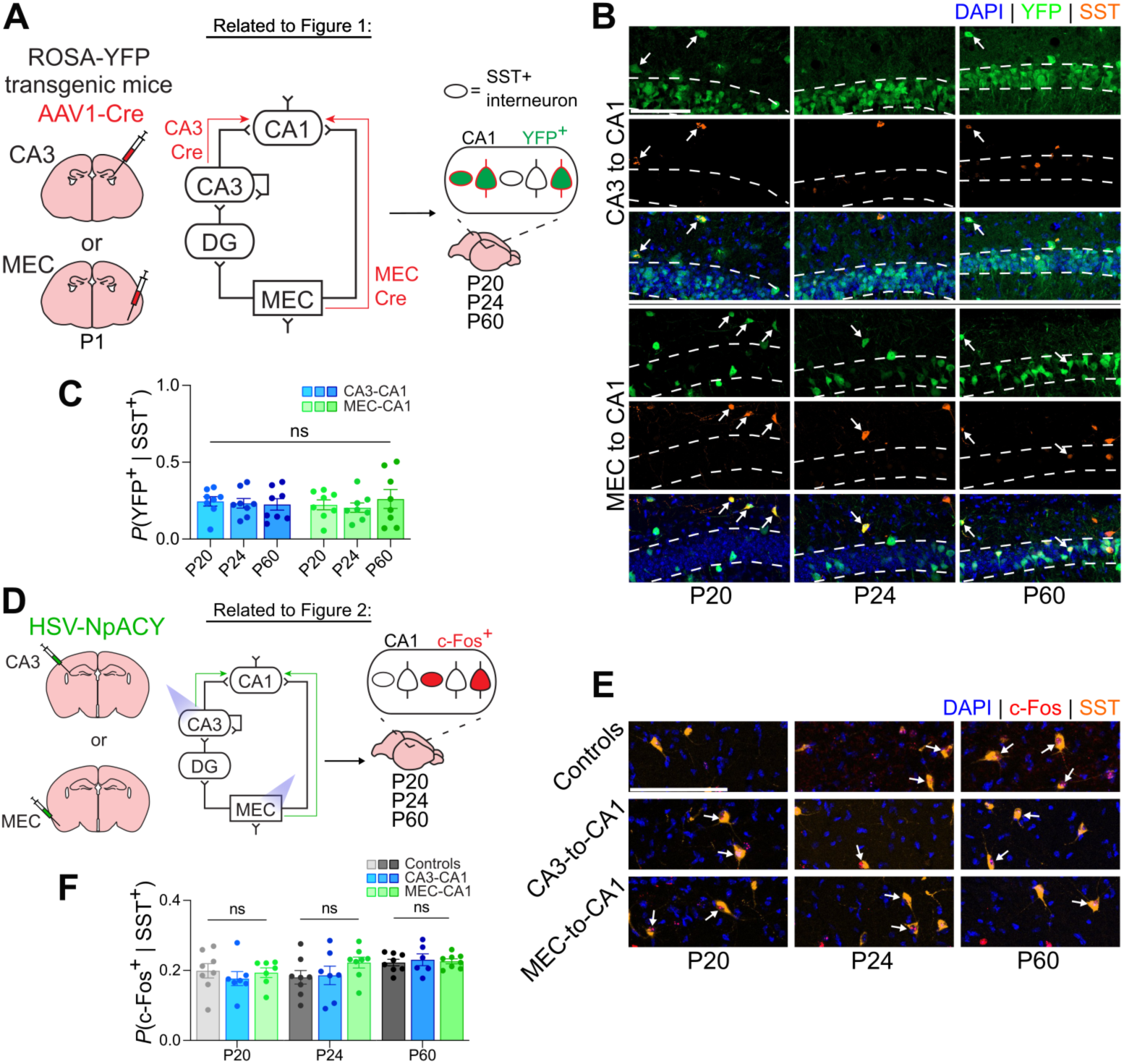
SST interneurons do not receive preferential input or activation from CA3 and MEC (Related to Figure 1 and Figure 2). **(A and B)** Quantification of transsynaptic labeling of somatostatin (SST^+^) CA1 interneurons following AAV1-Cre injection into CA3 or MEC. Arrows indicate co-labeled cells. **(C)** SST^+^ interneuron labeling did not show a developmental increase in CA3 or MEC projections to CA1 (ANOVA, age × projection: *F*_2,42_ = 0.42, *P* = 0.6631). **(D and E)** c-Fos expression in CA1 SST^+^ interneurons following photostimulation of CA3 or MEC at P20, P24, and P60. Arrows indicate co-labeled cells. **(F)** Neither CA3 nor MEC stimulation significantly increased c-Fos expression in SST^+^ interneurons across development (ANOVA, age × photostimulation: *F*_4,58_ = 0.76, *P* = 0.5567). Data points are individual mice with mean ± s.e.m. Scale bars: white = 100 µm. \**P* < 0.05; ** *P* < 0.01; \*\*\**P* < 0.001; \*\*\*\**P* < 0.0001.

**Fig. S4.**
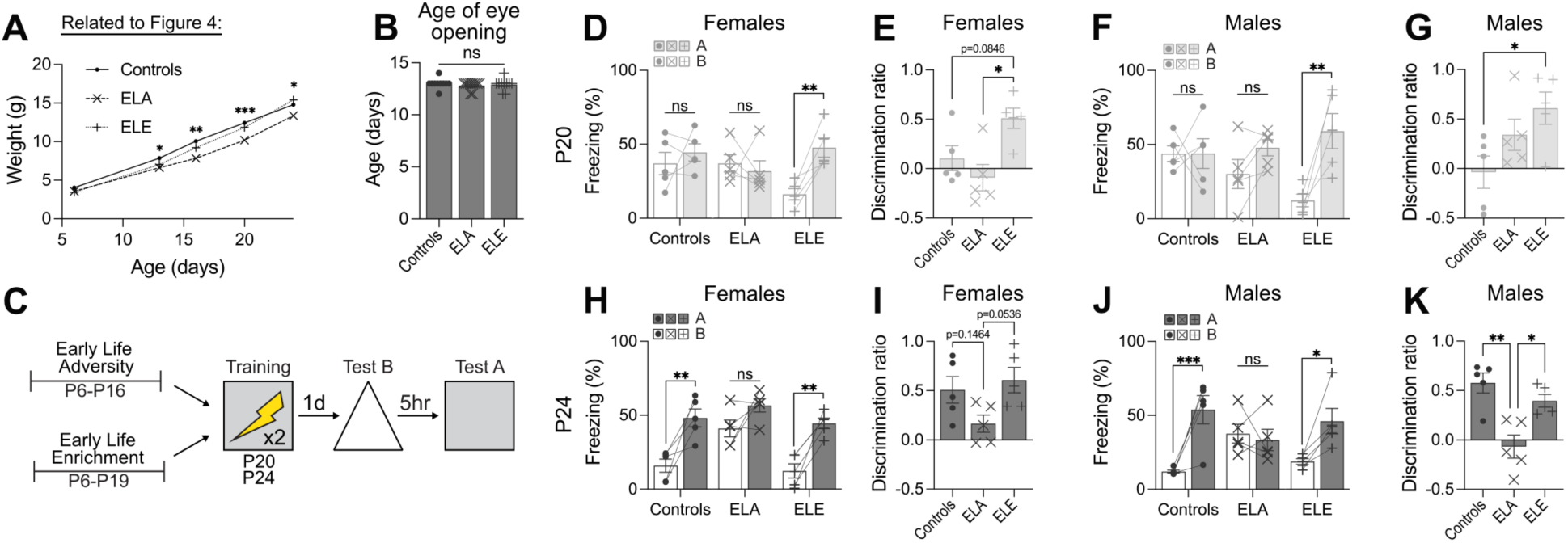
Early-life experience does not alter eye opening and produces comparable effects on memory resolution in males and females (Related to Figure 4). **(A and B)** ELA, but not ELE, reduced developmental weight gain (ANOVA, experience: *F*_2,27_ = 29.34, *P* < 0.0001), and neither manipulation altered the timing of eye opening (ANOVA, experience: *F*_2,27_ = 0.42, *P* = 0.6642). **(C)** Contextual fear memory following ELE or ELA in male and female mice. **(D and E)** ELE in P20 females resulted in context specific freezing (ANOVA, context × experience: *F*_2,12_ = 3.98, *P* = 0.0471) and high discrimination (ANOVA, experience: *F*_2,12_ = 6.39, *P* = 0.0129). **(F and G)** ELE in P20 males resulted in a trend towards context specific freezing (ANOVA, context × experience: *F*_2,12_ = 3.54, *P* = 0.0621) and high discrimination (ANOVA, experience: *F*_2,12_ = 4.06, *P* = 0.0451). **(H and I)** ELA in P24 females resulted in higher freezing (ANOVA, experience: *F*_2,12_ = 11.88, *P* = 0.0014) and a trend towards low discrimination (ANOVA, experience: *F*_2,12_ = 3.80, *P* = 0.0527). **(J and K)** ELA in P24 males resulted in context generalization (ANOVA, context × experience: *F*_2,12_ = 6.81, *P* = 0.0106) and low discrimination (ANOVA, experience: *F*_2,12_ = 12.00, *P* = 0.0014). Data points are individual mice with mean ± s.e.m. \**P* < 0.05; ** *P* < 0.01; \*\*\**P* < 0.001; \*\*\*\**P* < 0.0001.

**Fig. S5.**
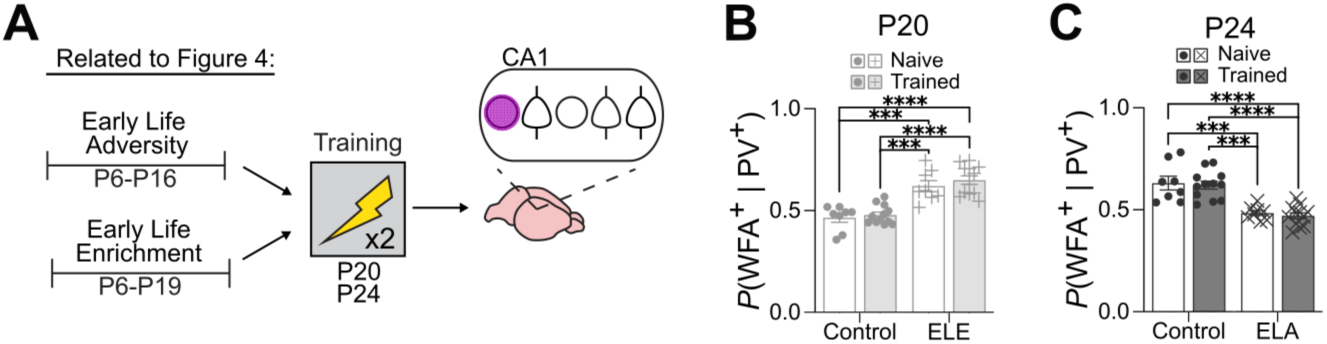
Early-life experience bidirectionally regulates PNN assembly around CA1 PV interneurons (Related to Figure 4). (**A)** Quantification of WFA^+^ perineuronal nets surrounding CA1 PV**^+^**interneurons following ELE or ELA and contextual fear training. **(B and C)** ELE increased PNN assembly around CA1 PV**^+^** interneurons at P20 in both naive and trained mice (ANOVA, experience: *F*_1,36_ = 60.69, *P* < 0.0001; training × experience: *F_1_*_,36_ = 0.13, *P* = 0.7182), whereas ELA reduced PNN assembly at P24 (ANOVA, experience: *F*_1,36_ = 51.42, *P* < 0.0001; training × experience: *F*_1,36_ = 0.01, *P* = 0.9100). Data points are individual mice with mean ± s.e.m. \**P* < 0.05; ** *P* < 0.01; \*\*\**P* < 0.001; \*\*\*\**P* < 0.0001.

**Fig. S6.**
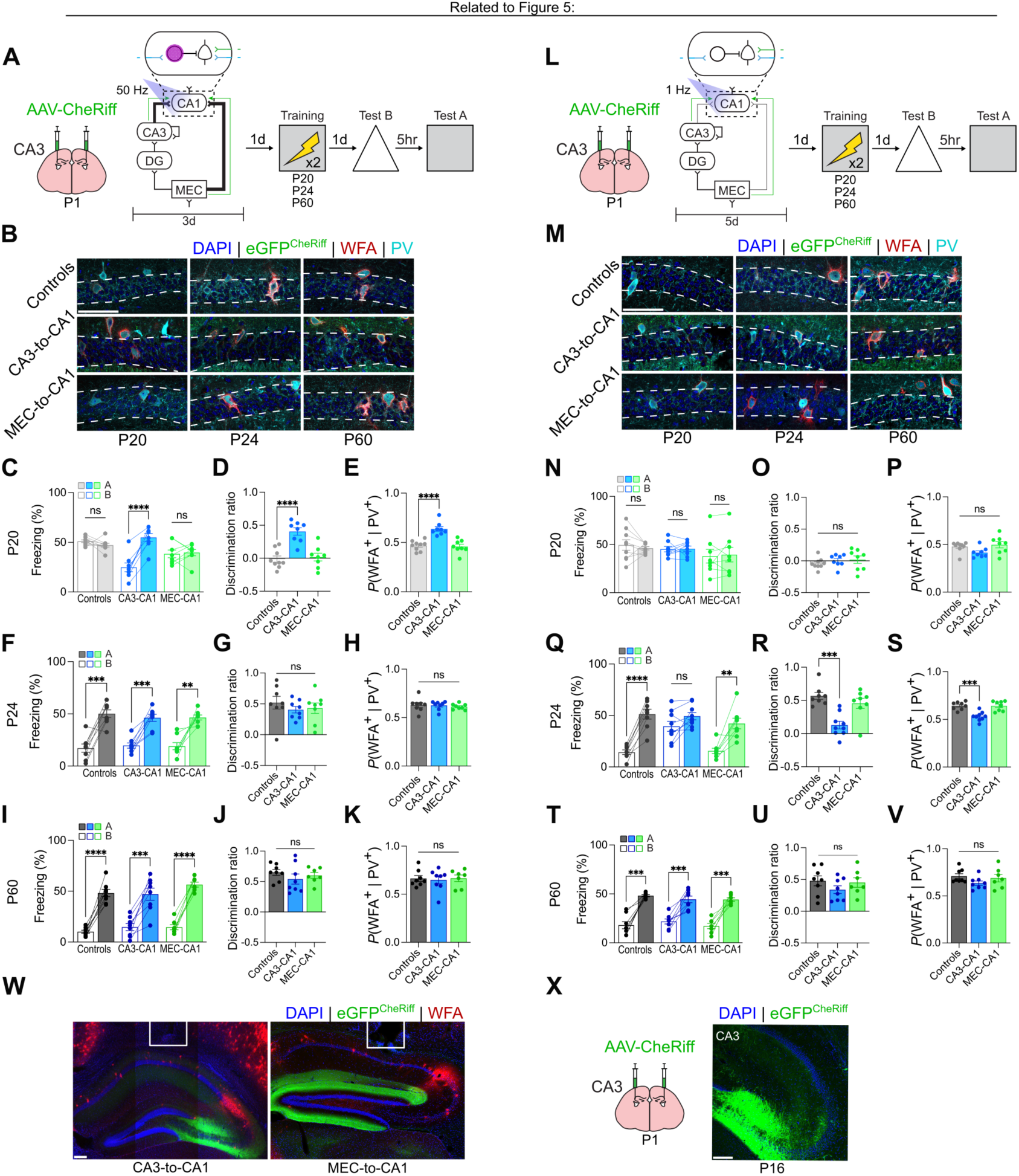
TSP activity during development regulates memory resolution and PNNs maturation in CA1 (Related to Figure 5). **(A and B)** Experimental design and quantification of PNN assembly following high-frequency photostimulation of CA3→CA1 and MEC→CA1 terminals during development. **(C and D)** High-frequency stimulation of CA3→CA1 terminals during P17–19 resulted in context specific freezing when mice were trained at P20, whereas stimulation of MEC→CA1 terminals did not alter freezing (ANOVA, photostimulation × context: *F*_2,22_ = 7.59, *P* = 0.0031) and discrimination (ANOVA, photostimulation: *F*_2,22_ = 18.66, *P* < 0.0001). **(E)** High-frequency stimulation of CA3→CA1 increased PNN assembly around CA1 PV^+^ interneurons at P20 (ANOVA, photostimulation: *F*_2,22_ = 21.84, *P* < 0.0001). **(F and G)** High-frequency stimulation during P21–23 did not alter freezing (ANOVA, photostimulation × context: *F*_2,21_ = 0.48, *P* = 0.6275) or discrimination (ANOVA, photostimulation: *F*_2,21_ = 0.52, *P* = 0.3365) when mice were trained at P24. **(H)** High-frequency stimulation did not affect PNN assembly around CA1 PV^+^ interneurons at P24 (ANOVA, photostimulation: *F*_2,21_ = 0.24, *P* = 0.7916). **(I and J)** High-frequency stimulation during P57–59 did not alter freezing (ANOVA, photostimulation × context: *F*_2,20_ = 1.13, *P* = 0.3419) or discrimination (ANOVA, photostimulation: *F*_2,20_ = 0.78, *P* = 0.4719) when mice were trained at P60. **(K)** High-frequency stimulation did not affect PNN assembly around CA1 PV^+^ interneurons at P60 (ANOVA, photostimulation: *F*_2,20_ = 0.07, *P* = 0.9323). **(L and M)** Experimental design and quantification of PNN assembly following low-frequency stimulation of CA3→CA1 and MEC→CA1 terminals during development. **(N and O)** Low-frequency stimulation during P17–19 did not alter freezing (ANOVA, photostimulation × context: *F*_2,22_ = 0.72, *P* = 0.4979) or discrimination (ANOVA, photostimulation: *F*_2,22_ = 0.61, *P* = 0.5502) when mice were trained at P20. **(P)** Low-frequency stimulation did not affect PNN assembly around CA1 PV^+^ interneurons at P20 (ANOVA, photostimulation: *F*_2,22_ = 3.28, *P* = 0.0567). **(Q and R)** Low-frequency stimulation of CA3→CA1 terminals during P21–23 resulted in context generalization when mice were trained at P24, whereas stimulation of MEC→CA1 terminals did not alter freezing (ANOVA, photostimulation × context: *F*_2,22_ = 8.67, *P* = 0.0017) and discrimination (ANOVA, photostimulation: *F*_2,22_ = 13.75, *P* = 0.0001). **(S)** Low-frequency stimulation of CA3→CA1 decreased PNN assembly around CA1 PV^+^ interneurons at P24 (ANOVA, photostimulation: *F*_2,22_ = 16.46, *P* < 0.0001). **(T and U)** Low-frequency stimulation during P57–59 did not alter freezing (ANOVA, photostimulation × context: *F*_2,20_ = 0.96, *P* = 0.3986) or discrimination (ANOVA, photostimulation: *F*_2,20_ = 1.11, *P* = 0.3486) when mice were trained at P60. **(V)** Low-frequency stimulation did not affect PNN assembly around CA1 PV^+^ interneurons at P60 (ANOVA, photostimulation: *F*_2,20_ = 1.69, *P* = 0.2100). **(W)** Representative images of AAV-CheRiff expression and PNNs labelled with WFA and optic fibers above CA1. **(X)** AAV-CheRiff was strongly expressed by P16 following injection at P1. Data points are individual mice with mean ± s.e.m. Scale bars: white = 100 µm. \**P* < 0.05; ** *P* < 0.01; \*\*\**P* < 0.001; \*\*\*\**P* < 0.0001.

**Fig. S7.**
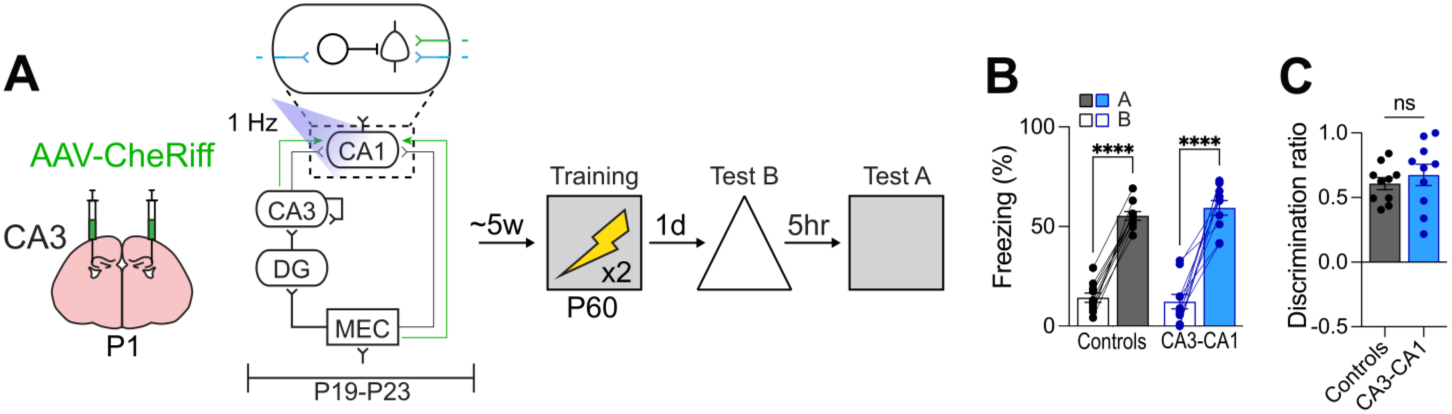
Effects of reduced TSP activity during development on memory resolution are transient. **(A)** Experimental design for assessing the persistence of the effects of low-frequency stimulation of CA3→CA1 terminals. Mice received low-frequency stimulation on P19–23 and were subsequently trained in contextual fear conditioning at P60. **(B and C)** Mice exhibited context specific freezing (ANOVA, photostimulation × context: *F*_1,18_ = 0.95, *P* = 0.3431) and high discrimination (unpaired *t*-test, *t*_18_ = 0.71, *P* = 0.4869) when tested at P60. Data points are individual mice with mean ± s.e.m. \**P* < 0.05; ** *P* < 0.01; \*\*\**P* < 0.001; \*\*\*\**P* < 0.0001.

**Fig. S8.**
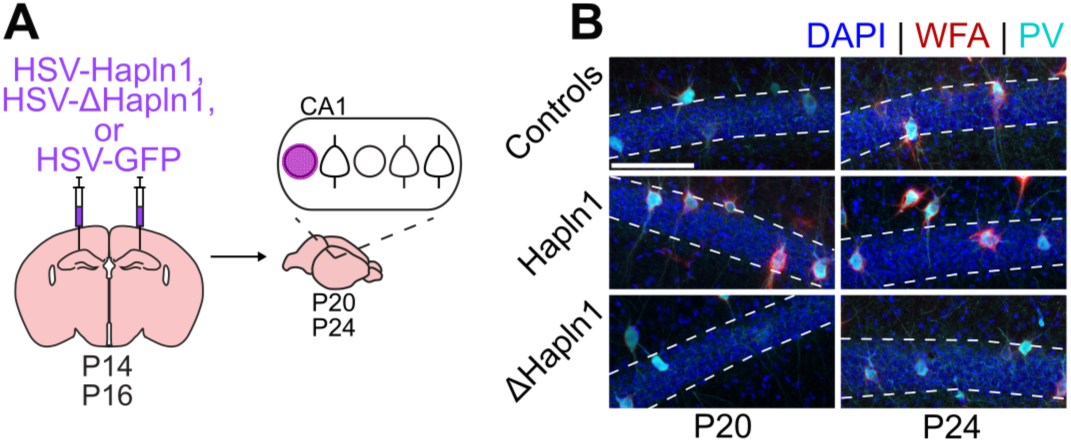
Viral manipulations targeting HAPLN1 modulate PNN maturation in CA1. **(A and B)** HSV-Hapln1 increased PNNs in CA1 at P20 when injected at P14. HSV-ΔHapln1 reduced PNNs in CA1 at P24 when injected at P16. Scale bars: white = 100 µm.

## Notes

### Competing Interest Statement

The authors have declared no competing interest.

## References and Notes

1. J. Piaget, B. Inhelder, Memory and Intelligence (Basic Books, 1973).

2. C. T. Ellis et al., Current Biology. 31, 3358–3364.e4 (2021).

3. J. R. Saffran, N. Z. Kirkham, Annu. Rev. Psychol. 69, 181–203 (2018).

4. C. O’Doherty et al., Nat Neurosci. 29, 693–702 (2026).

5. H. K. Lambert et al., J. Neurosci. 37, 1925–1934 (2017).

6. J. Bevandić et al., Neuron. 112, 1060–1080 (2024).

7. P. J. Bauer, Developmental Psychobiology. 50, 19–31 (2008).

8. C. T. Ngo, N. S. Newcombe, I. R. Olson, Dev Sci. 21, e12556 (2017).

9. L. Rollins, E. B. Cloude, Learn. Mem. 25, 294–297 (2018).

10. A. Keresztes, C. T. Ngo, U. Lindenberger, M. Werkle-Bergner, N. S. Newcombe, Trends in Cognitive Sciences. 22, 676–686 (2018).

11. A. I. Ramsaran et al., Science. 580, 543–551 (2023).

12. A. I. Ramsaran, M. L. Schlichting, P. W. Frankland, Developmental Cognitive Neuroscience. 36, 1–15 (2019).

13. M. Lahr, F. Imhof, L. Mauro, T. Ulmer, F. Donato, BioRxiv (2025), doi:10.1101/2025.09.27.678956.

14. M. J. Anderson, D. C. Riccio, Learning & Behavior. 33, 444–453 (2005).

15. J. Bevandić, F. Stella, H. F. Ólafsdóttir, Current Biology. 34, 5062–5072.e5 (2024).

16. A. C. Schapiro, N. B. Turk-Browne, M. M. Botvinick, K. A. Norman, Phil. Trans. R. Soc. B. 372, 20160049 (2017).

17. A. Bakker, C. B. Kirwan, M. Miller, C. E. L. Stark, Science. 319, 1640–1642 (2008).

18. S. Baker et al., Current Biology. 26, 2629–2634 (2016).

19. M. A. Yassa, C. E. L. Stark, Trends in Neurosciences. 34, 515–525 (2011).

20. T. Kitamura et al., Neuron. 87, 1317–1331 (2015).

21. J. Sucevic, A. C. Schapiro, BioRxiv (2022), doi:10.1101/2022.01.12.476051.

22. M. Gumus, M. L. Mack, Proc. Natl. Acad. Sci. U.S.A. 123, e2503388123 (2026).

23. D. Singh, A. C. Schapiro, Philosophical Transactions of the Royal Society B: Biological Sciences. 381 (2026), doi:10.1098/rstb.2025.0243.

24. C. McKenzie, et al., BioRxiv (2026).

25. E. Atucha, S.-P. Ku, M. T. Lippert, M. M. Sauvage, Cell Reports. 42, 113317 (2023).

26. R. L. Gómez, J. O. Edgin, Developmental Cognitive Neuroscience. 18, 57–69 (2016).

27. P. Lavenex, P. Banta Lavenex, Behavioural Brain Research. 254, 8–21 (2013).

28. R. Cossart, R. Khazipov, Physiological Reviews (2021), doi:10.1152/physrev.00044.2020.

29. F. Donato et al., The Journal of Neuroscience. 41, 920–926 (2021).

30. H. F. Ólafsdóttir, D. Bush, C. Barry, Current Biology. 28, R37–R50 (2018).

31. A. I. Ramsaran et al., Current Biology (2024), doi:10.1101/2024.11.06.622296.

32. M. Hernández-Frausto, O. M. Bilash, A. V. Masurkar, J. Basu, Front. Neural Circuits. 17, 1223891 (2023).

33. A. Khlaifia, E. Honoré, J. Artinian, I. Laplante, J.-C. Lacaille, Mol Brain. 15, 56 (2022).

34. R. Zemankovics, J. M. Veres, I. Oren, N. Hajos, Journal of Neuroscience. 33, 12337–12351 (2013).

35. H. Hu, J. Gan, P. Jonas, Science. 345, 1255263 (2014).

36. A. Travaglia, R. Bisaz, E. S. Sweet, R. D. Blitzer, C. M. Alberini, Nat Neurosci. 19, 1225– 1233 (2016).

37. S. N. Duffy, K. J. Craddock, T. Abel, P. V. Nguyen, Learn. Mem. 8, 26–34 (2001).

38. L. Malave, M. T. Van Dijk, C. Anacker, Transl Psychiatry. 12, 306 (2022).

39. F. Donato, R. I. Jacobsen, M.-B. Moser, E. I. Moser, Science. 355, eaai8178 (2017).

40. L. C. Katz, C. J. Shatz, Science. 274, 1133–1138 (1996).

41. A. K. Short, T. Z. Baram, Nat Rev Neurol. 15, 657–669 (2019).

42. B. E. Sherman, N. B. Turk-Browne, E. V. Goldfarb, Perspect Psychol Sci. 19, 103–125 (2024).

43. C. J. Brainerd, V. F. Reyna, Developmental Review. 24, 396–439 (2004).

44. E. Favuzzi et al., Neuron. 95, 639–655.e10 (2017).

45. A. Dityatev et al., Developmental Neurobiology. 67, 570–588 (2007).

46. T. K. Hensch, Annual Review of Neuroscience. 27, 549–579 (2004).

47. E. Dzyubenko et al., Cell. Mol. Life Sci. 78, 5647–5663 (2021).

48. J. T. Sakata et al., J. Neurosci. 45, e1367252025 (2025).

49. C. M. Alberini, A. Travaglia, J. Neurosci. 37, 5783–5795 (2017).

50. M. T. Birnie, T. Z. Baram, Science. 376, 1055–1056 (2022).

51. D. M. Barch, M. P. Harms, R. Tillman, E. Hawkey, J. L. Luby, Journal of Abnormal Psychology. 128, 81–95 (2019).

52. A. L. Decker, K. Duncan, A. S. Finn, D. J. Mabbott, Nat Commun. 11, 4040 (2020).

## References only cited in the supplementary materials

53. S. Srinivas et al., BMC Dev Biol. 1, 4 (2001).

54. S. Hippenmeyer et al., PLoS Biol. 3, e159 (2005).

55. A. J. Rashid et al., Science. 353, 383–387 (2016).

56. S. J. Russo et al., J. Neurosci. 29, 3529–3537 (2009).

57. B. Zingg et al., Neuron. 93, 33–47 (2017).

58. B. Zingg, B. Peng, J. Huang, H. W. Tao, L. I. Zhang, J. Neurosci. 40, 3250–3267 (2020).

59. D. R. Hochbaum et al., Nat Methods. 11, 825–833 (2014).

60. J. J. Kim, M. S. Fanselow, Science. 256, 675–677 (1992).

61. S. Y. Ko et al., Nature. 643, 735–743 (2025).

